# *In vivo* macromolecular crowding is differentially modulated by Aquaporin 0 in zebrafish lens: insights from a nano-environment sensor and spectral imaging

**DOI:** 10.1101/2021.05.15.442187

**Authors:** Irene Vorontsova, Alexander Vallmitjana, Belén Torrado, Thomas Schilling, James E. Hall, Enrico Gratton, Leonel Malacrida

## Abstract

Macromolecular crowding is crucial for cellular homeostasis. *In vivo* studies of macromolecular crowding and ultimately water-dynamics are needed to understand their role in cellular fates. The macromolecular crowding in the lens is essential for understanding normal optics of the lens, and moreover for understanding and prevention of cataract and presbyopia. Here we combine the use of the water nano-environmentally sensitive sensor (6-acetyl-2-dimethylaminonaphthalene, ACDAN) with *in vivo* studies of Aquaporin zero zebrafish mutants to understand the lens macromolecular crowding. Spectral phasor analysis of ACDAN fluorescence reveal the extent of water dipolar relaxation and demonstrate that the mutations in the duplicated zebrafish Aquaporin 0s, Aqp0a and Aqp0b, alter the water state and macromolecular crowding in the living zebrafish lens. Our results provide *in vivo* evidence that Aqp0a promotes fluid influx in the deeper lens cortex, whereas Aqp0b facilitates fluid efflux. This work opens new perspectives for *in vivo* studies on macromolecular crowding.

**Teaser:** In this study we uncover the roles of Aquaporin 0 in macromolecular crowding required for lens development and vision.

## Introduction

Solutes occupy 35-95% of eukaryotic cell volume, with the rest being water. The ratio and interaction between water and solutes determine cellular macromolecular crowding, a key feature of cellular organization and function (1–3). However, its physiological roles remain obscure and are challenging to study in living organisms. There has been a great effort to understand the roles of macromolecular crowding in enzymatic activity, cell physiology, and pathophysiology, primarily using *in vitro* and cell culture systems (1, 3– 5). However, efforts to study macromolecular crowding *in vivo* with non-invasive, high-resolution spectroscopic tools remain challenging. In this study, we were able to overcome the technical difficulties to study the *in vivo* development of macromolecular crowding in the lens, as well as to test the contribution of the zebrafish Aquaporin 0 orthologues to macromolecular crowding by using a nano-environmental sensor (ACDAN, 6-acetyl-2-dimethylaminonaphthalene) paired with hyperspectral imaging and spectral phasor analysis.

The ocular lens relies on high macromolecular crowding to determine its structure and function (6). To achieve and maintain the required refractive index gradient the lens fiber cells are enriched with crystallin proteins (4), moreover, lens water is tightly regulated to maintain lens homeostasis and proper optics. Regulation of water transport/activity is finely tuned in different parts of the lens, adjusting macromolecular crowding to optimize the refractive index gradient (7). Water influx/efflux is required to facilitate the microcirculation of ions by inward fluid flow at the lens poles and efflux of nutrients and waste at the equator (8). In addition, the direction of fluid and ion transport reverse with depth into the lens due to changes in electrochemical gradients (9). Net fluid influx in the inner lens cortex must equal net fluid efflux in the outer cortex to maintain homeostasis (10). The mechanisms by which water transport regulates macromolecular crowding in the lens *in vivo* remain unclear; nonetheless, Aquaporin 0 (AQP0, also known as Membrane Intrinsic Protein, MIP), and in mammals Aquaporin 5, were proposed as water transport regulators (11, 12). AQP0 is the most abundant membrane protein in the lens and is required for lens homeostasis (6). AQP0 permeates water *in vitro* (13) and functions as an adhesive protein (14, 15), cytoskeletal anchor (16, 17), and regulator of gap junctions in lens fiber cells (18, 19). In mammals, a single AQP0 protein performs all of these functions. In contrast, zebrafish (*Danio rerio*), as a consequence of an ancient teleost-lineage genome duplication, have two AQP0 orthologues (20), Aqp0a and Aqp0b, allowing genetic dissection of at least some of these functions (21–23). Aqp0a and Aqp0b are both essential for lens transparency at 3 days post fertilization (dpf) (21–23). While the loss of Aqp0b alone causes no apparent lens defects, loss of Aqp0a disrupts the anterior lens suture, leading to anterior polar opacity at adult stages (21, 24). Both Aqp0a and Aqp0b expressed in *Xenopus laevis* ooctyes, permeate water *in vitro* (22, 25). However, only Aqp0a knock-down zebrafish required introduction of Aqp0 with intact water transport function to rescue embryonic cataract (23), possibly by recovering macromolecular crowding homeostasis.

Measuring water homeostasis in living lenses in the presence or absence of Aqp0 function in specific lens regions is challenging due to the dearth of non-invasive tools. Our approach to investigating water homeostasis *in vivo* employs a solvatochromic molecule (ACDAN), and hyperspectral imaging to provide information on spectroscopic macromolecular crowding at the nano-environment sensor. ACDAN is a non-toxic membrane permeable fluorescent probe, which reports on water dipolar relaxation (DR) within a few Angstroms of its location (26, 27). This probe has been used as a sensor for water dynamics in living cells (28, 29). Water dipolar relaxation (DR) is responsive to macromolecular crowding due to its extreme sensitivity to the number of water molecules and the dipolar relaxation time in its nano-environment (see schematic at Figure 1A). In this study, we define a dipolar relaxation index using hyperspectral microscopy and spectral phasor analysis. Changes in ACDAN DR index are indicative of macromolecular crowding variation. This index was built using spectral phasor plot analysis (30, 31). The spectral phasor transforms spectra pixel-by-pixel into a 2D scatter plot, where the axes are the real and imaginary components of the Fourier transform. This transformation captures the morphology of the emission spectra and maps it onto the phasor 2D space where, in polar coordinates, the angle carries the information regarding the spectral center of mass, and the radial direction carries information on the spectral broadening. The spectral phasor properties are crucial for the pixel-by-pixel model-less analysis of the spectral information (32), as described in the methods section.

**Figure 1:**
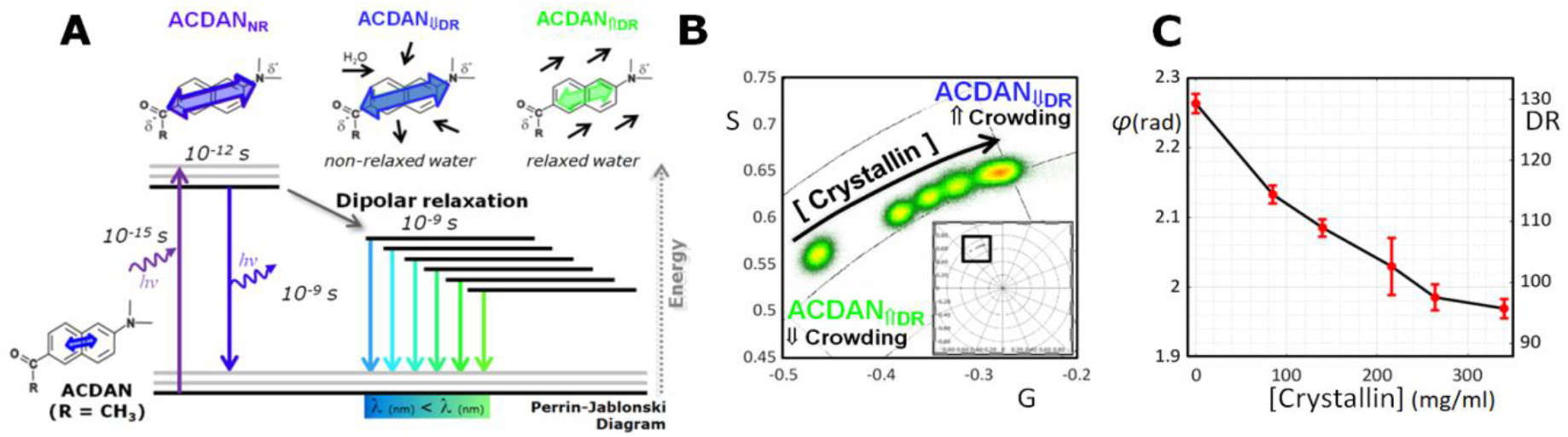
Solvatochromic properties of ACDAN measure water homeostasis and macromolecular crowding. **(A)** ACDAN photophysics display strong sensitivity to the polarity of the environment and solvent relaxation. In an uncrowded environment high dipolar relaxation (DR) results in a spectral red-shift, while in crowded environments low DR results in a spectral blue-shift. **(B)** Antarctic tooth fish γM8d crystallin at concentrations of 0, 85, 140, 216, 264, and 340 mg/ml was imaged with 5 µM ACDAN *in vitro*. A blue spectral shift was observed in response to increased macromolecular crowding, and thus a decrease in DR. **(C)** The mean phasor phase angle/DR show an inverse relationship with crystallin concentration (n=3). Notice the DR scale was extended over 100% to include the spectral shift found at the crystallin solutions, while lens DR was in the 0 to 100 DR.

To tackle the spatial/temporal DR information in the lens, we developed a combination of hyperspectral imaging and image processing tools for 3D (x/y/λ) to 5D (x/y/z/λ/t) analysis to non-invasively study the development of lens macromolecular crowding and its perturbation in zebrafish Aqp0 mutant lenses. In this work, ACDAN’s DR index was first shown to be sensitive to crystallin crowding *in vitro* and macromolecular crowding in living zebrafish lenses, validating its utility for measuring water dynamics *in vivo*.

ACDAN imaging in embryonic and larval lenses reveals a differential increase in macromolecular crowding throughout development, in a spatial and temporal concert. The macromolecular crowding revealed by ACDAN within the lens was higher than it could achieve in *in vitro* crystallin samples. We show that Aqp0a is required for the normal development of macromolecular crowding in the lens cortex, particularly at the posterior pole. Putting these data in light with previous work, we show that Aqp0a facilitates water influx in the inner cortex and efflux in the outer lens cortex. While the function of Aqp0b is non-essential, in combination with the loss of Aqp0a, it results in extensive cell swelling in the cortex, suggesting Aqp0b’s role is water efflux in the outer cortex. We have thus dissected the specific roles for Aqp0a and Aqp0b in establishing and maintaining regional lens macromolecular crowding.

## Results

### Spectral imaging of ACDAN reveals macromolecular crowding in vitro

ACDAN spectral imaging provides a read-out of environment DR within a few angstroms of its macromolecular environment (Figure 1A). A solution of ACDAN in water has a peak fluorescence emission in the green region of the spectrum, around 520nm, which shifts towards blue (shorter wavelengths) with an increasing protein-to-water ratio (Figure 1B). In the excited-state, ACDAN has a higher dipole moment than the ground state, and movement of nearby dipole-active molecules, such as water, lead to dipole relaxation (DR) (33), observed as a shift towards longer wavelengths (red-shift) in the maximum of the fluorescence emission. To measure these wavelength shifts in the emission, we used hyperspectral imaging, collecting the entire ACDAN spectrum and then transform to phasor space to map the pixels onto the spectral phasor plot (Figure 1B) (32). The phasor transform’s power relies on its fit-free approach, meaning *apriori* knowledge of the spectral emission characteristics is not required.

To test spectral emission responses of ACDAN to increasingly crowded environments related to the intracellular lens environment, we imaged a range of Antarctic tooth fish γM8d crystallin concentrations in solution. Consistent with ACDAN serving as an accurate quantitative read-out of the water activity (confinement), and therefore protein-water ratio, we observed a blue spectral emission shift (DR decrease) as crystallin concentration increased (Figure 1B), confirming its sensitivity to macromolecular crowding in a lipid-free environment. Figure 1C shows the delta phase due to the decrease in DR when the concentration of γM8d crystallin increases from 0 to 340 mg/ml, which is close to maximum crystallin solubility. Note, that ACDAN short shifts in the delta phase represent significant increases in the molecular crowding.

### Macromolecular crowding increases during lens development

We developed an automated image processing pipeline to analyze the hyperspectral stack in the zebrafish lens (Figure 2A). The details of this experimental pipeline can be found in Supplementary Figure 3. Spatial DR is crucial for understanding water dynamics in the lens, so the ACDAN spectra were transformed pixel-by-pixel into the spectral phasor plot (Figure 2B), and then following the reciprocity principle, the DR value for each pixel (color scale) was applied back to its original x-y location (Figure 2C). To further analyze regional lens DR, images were segmented into different lens regions, measuring the distribution of DR across the radial geometry of the lens (Figure 2D, Supplementary Figure 2), and z-stacks were carried out. This analysis is fundamental to further understand the spatial-temporal information during the lens development and growth, for instance, 3D geometry (Supplementary Figure 4) and the axial DR (Figure 7) (cover in the following sections).

**Figure 2:**
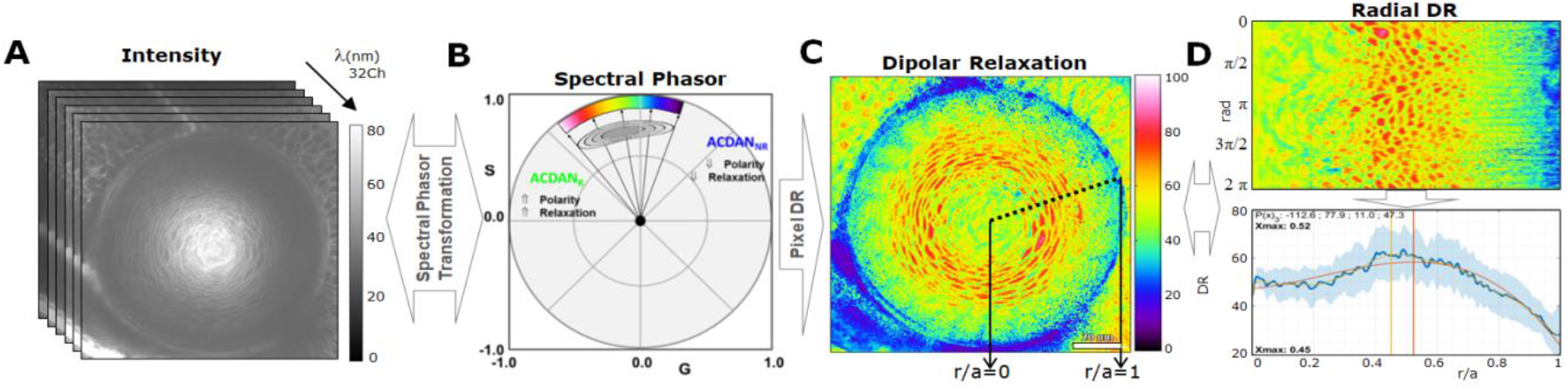
Characterization of lens ACDAN emission using the phasor approach. **(A)** Hyperspectral images were transformed to the spectral phasor plot **(B)**, which separates the signals from relaxed and unrelaxed dipolar states. **(C)** Dipolar relaxation (DR) values were applied back to the original image pixel-by-pixel, which were then processed for parameter extraction. **(D)** Radial analysis of mean DR signal from the center of the lens (r/a = 0, where r = distance from lens center, a = lens radius) to lens periphery (r/a=1) is shown. The polar geometry is transformed to cartesian geometry, the horizontal direction being the radius, vertical angle, and the mean DR value is then graphed (bottom panel), enabling analysis of regional change of mean DR. A polynomial fit was used to estimate max DR. Details can be found in the Supplementary Figure 2.

Crystallins are by far the most abundant cytoplasmic lens proteins. They become more concentrated toward the lens center, generating the gradient of refractive index required for emmetropia. We visualized the DR during the formation of this gradient in the zebrafish lens *in vivo* using ACDAN. The ACDAN signal was analyzed in embryonic and larval lenses in the equatorial plane. Since fish grow at different rates depending on their environment, lens diameters as well as age were used as measures of development during analysis (Supplementary Figure 5A) as we previously used (34).

ACDAN intensity images do not provide information regarding water dynamics; however, when transformed to DR images the spatial information across the lenses is valuable (Supplementary Figure 6). One may notice that the phasor cloud obtained for the different stages (Supplementary Figure 6 B1-6) are blue-shifted (lower phase) with respect to the higher crystallin concentration (340 mg/ml) obtained *in vitro* (Figure 1B and D). This result indicates that higher macromolecular crowding is experienced by ACDAN within lens cells.

In the immature lens at 2 days post fertilization (dpf) a high DR signal was found at the lens core, as well as in cell nuclei in the lens cortex and nucleus (Figure 3A1). Higher DR in the cell nuclei compared to the cytoplasm has been previously observed in cell culture (data not shown) and indicates lower nuclear macromolecular crowding. Although by 3 dpf cell nuclei and organelles in fiber cells of the lens nucleus are degraded as part of the maturation of the lens (24, 35), the highest DR signal, which was in the cytoplasm of fiber cells, was still in the center of the lens, compared to lower DR at the lens periphery (Figure 3A2). As the lens matured, the lens nucleus gradually decreased in DR, and the maximum DR signal moved to the lens cortex, indicating an increased macromolecular crowding in the lens nucleus (Figure 3A3-6). The smooth mean radial profile confirmed the shift of the mean DR signal from the lens nucleus (r/a = 0) in young lenses to the cortex in older/larger lenses (Figure 3B). The maximum DR signal stabilized at a relative distance from the lens center of r/a ∼ 0.7 from around 5 dpf and older (Figure 3C). A sigmoid curve was used to model the transition between the two states and the fit to this curve was very high (R^2^=0.94). Regionally segmented DR showed a decrease in DR signal in the inner cortex (Figure 3D4) and more so in the lens nucleus with development (Figure 3D5).

**Figure 3:**
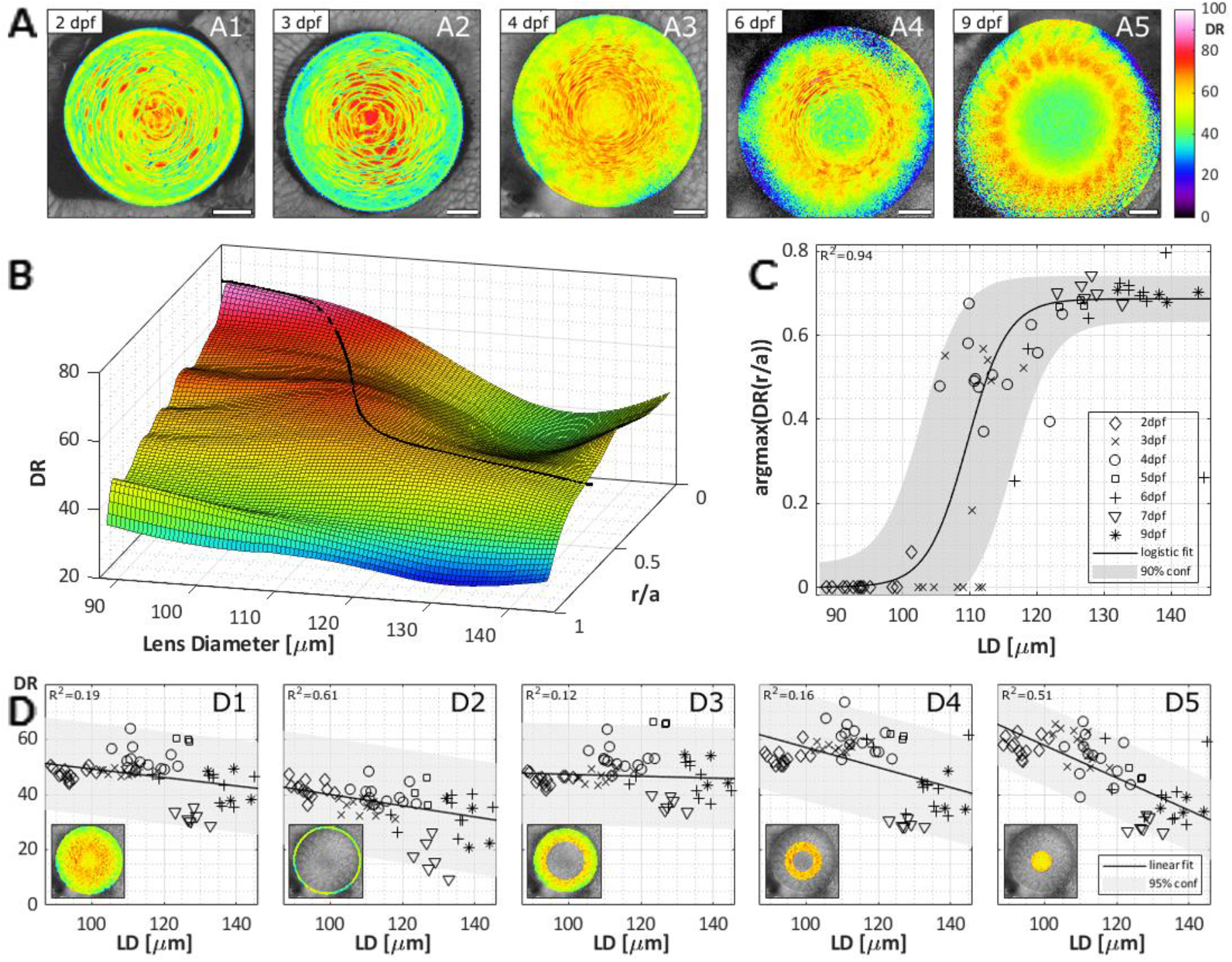
Distribution of dipolar relaxation in equatorial planes of zebrafish lenses during development. **(A)** Examples of dipolar relaxation (DR) images of lenses at specified days post fertilization (dpf). Scale bars are 20μm. **(B)** Smoothed surface of the mean DR radial profile (r/a) as a function of lens diameter (LD) (N=63). **(C)** Radial lens position of the maximal DR value with development. The data is fit to a sigmoid and in turn represented on the surface in panel B. **(D)** Mean DR of the whole equatorial lens plane **(D1)**, the epithelium **(D2)**, outer cortex **(D3)**, inner cortex **(D4)** and nucleus **(D5)** of the lens as indicated by the insets. See Supplementary Table 2 for a summary of n numbers. See Supplementary Figure 7 for ACDAN intensity images, spectral phasor plots, and DR images for examples shown in A.

Regions 5-10 *μ*m, and 75-85 *μ*m from the anterior pole (see Supplementary Table 1) were imaged to determine changes in DR as anterior and posterior sutures formed (Figure 4), respectively. DR decreased slightly in anterior sutures with lens development (Figure 4B4), while the lens epithelium (Figure 4B2) and cortex (Figure 4B3) DR were unchanged. The mean DR in anterior planes overall appears similar at all stages, while the max DR slightly shifts away from the deeper cortex with development (Supplementary Figure 7A-B). DR at posterior poles initially increased from 2 to 4 dpf, and then decreased in older lenses (Figure 4C-D). This trend was particularly apparent in the posterior suture (Figure 4D3). These trends are verified by a mean DR smoothed surface plot and maximum DR plot (Supplementary Figure 7C-D). Posterior regions had higher DR compared to the anterior regions (Supplementary Figure 8 A, E, I).

**Figure 4:**
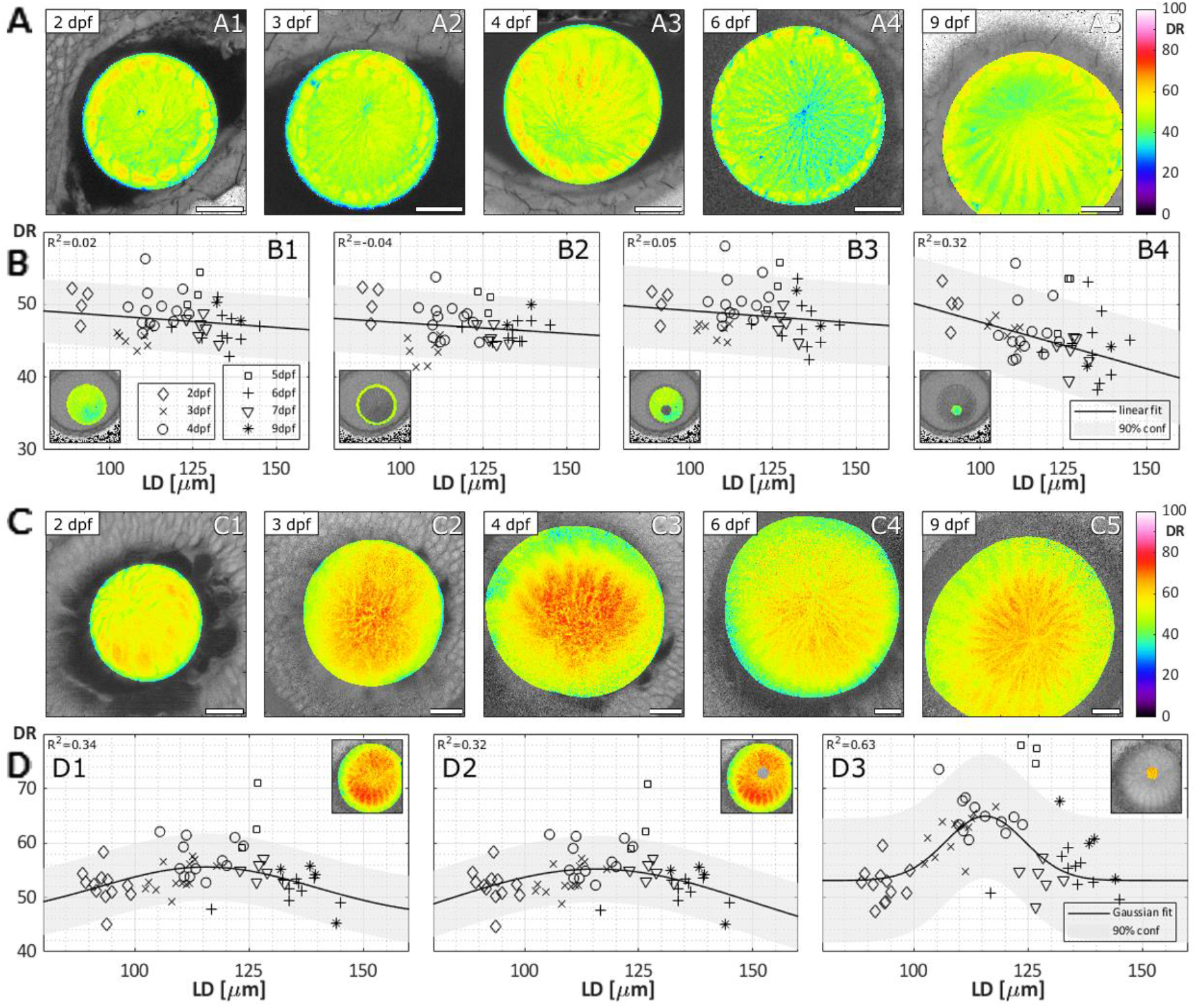
Distribution of dipolar relaxation in anterior and posterior planes of zebrafish lenses during development. **(A)** Examples of dipolar relaxation (DR) images of lenses obtained at the anterior pole at specified days post fertilization (dpf). Scale bars are 20μm. **(B)** Mean DR of the whole anterior lens plane **(B1)**, the epithelium **(B2)**, fiber cells **(B3)**, and sutural region **(B4)** as indicated by the insets (N=45). **(C)** Examples of DR images of lenses obtained at the posterior pole at specified dpf. Scale bars are 20 μm. **(D)** Mean DR of the whole posterior lens plane **(D1)**, fiber cells **(D2)**, and sutural region **(D3**; N=59). See Supplementary Table 2 for a summary of n numbers.

Imaging z-stacks of several lenses further confirmed these DR patterns. Examples of 2-4 dpf lens z-stacks in equatorial and axial orientations are provided as Supplementary animations 1-6. To confirm the DR patterns observed in mature lenses were not artifacts of the optical setup (e.g. low in the center, high at periphery), a spherically and conically slicing of the z-stacks was performed. Spherical slices were obtained as described for the equatorial, anterior and posterior planes by measuring the mean radial DR profile for each of the planes in the z-stack. We then obtained the DR distribution for each of these planes. The same procedure was performed in conical sections at different angles from the center of the lens to obtain the DR profile for each cone. By finding the maximum DR using the polynomial fit described previously and plotting it for the two spatial distributions, one can observe how the radial position of the maximum remains constant for the conical slices but not for the parallel slices, proving the geometry is spherical (Supplementary Figure 4). This analysis confirmed that the observed DR patterns were an intrinsic property of the lens and were not artifacts.

### Loss of Aqp0a function regionally disrupts macromolecular crowding in the lens

To understand the roles of Aqp0a and Aqp0b in the lens water homeostasis, and therefore macromolecular crowding, spatial studies of DR were carried out in null mutants previously generated by CRISPR-Cas9 gene editing (21). We focused on 4 dpf, as this stage follows loss of organelles from the lens nucleus and embryos have hatched, but the signal from the lens is still strong and not obscured by lens density and peripheral eye structures. Furthermore, our DR analysis during development showed that DR is lower in the nucleus at 4 dpf suggesting maturation of the macromolecular crowding and likely lens optics. See Supplementary Figure 9 for ACDAN intensity images to DR transformation data. Equatorial images of ACDAN emission revealed uniformly lower DR in *aqp0a*^*-/-*^ mutant lenses (Figure 5A2) compared with WT (Figure 5A1) and swollen cells with high DR, particularly in the cortex in double *aqp0a*^*-/-*^*/aqp0b*^*-/-*^ mutant lenses (Figure 5A4). Furthermore, double *aqp0a*^*-/-*^*/aqp0b*^*-/-*^ mutant lenses had increased diameters (Supplementary Figure 5B), while the other three genotypes were indistinguishable, indicating that the whole lens in double *aqp0a*^*-/-*^*/aqp0b*^*-/-*^mutants swelled. In contrast, DR levels and distribution in *aqp0b*^*-/-*^ mutants (Figure 5A3) appeared very similar to WT (Figure 5A1), with lower DR in the nucleus and elevated DR more peripherally. Mean DR values as a function of lens depth confirmed that *aqp0a*^*-/-*^ mutant lenses had a lower DR, particularly in the cortex, compared to other genotypes (Figure 5B, r/a>0.4). There was variability in phenotype penetrance in double *aqp0a*^*-/-*^*/aqp0b*^*-/-*^ mutant lenses (Supplementary Figure 10), as previously observed (21). However, double mutant lenses had higher mean DR in the outer cortex compared to WT and *aqp0a*^*-/-*^ mutant lenses (Figure 5B, r/a>0.6), and the maximum DR value was closer to the lens periphery (Figure 5C). Analyses of masked regions of the lens confirmed that the epithelium and cortex of *aqp0a*^*-/-*^ mutant lenses, but not the lens nucleus, had a lower DR than WT (Figure 5D) when measured at these equatorial planes.

**Figure 5:**
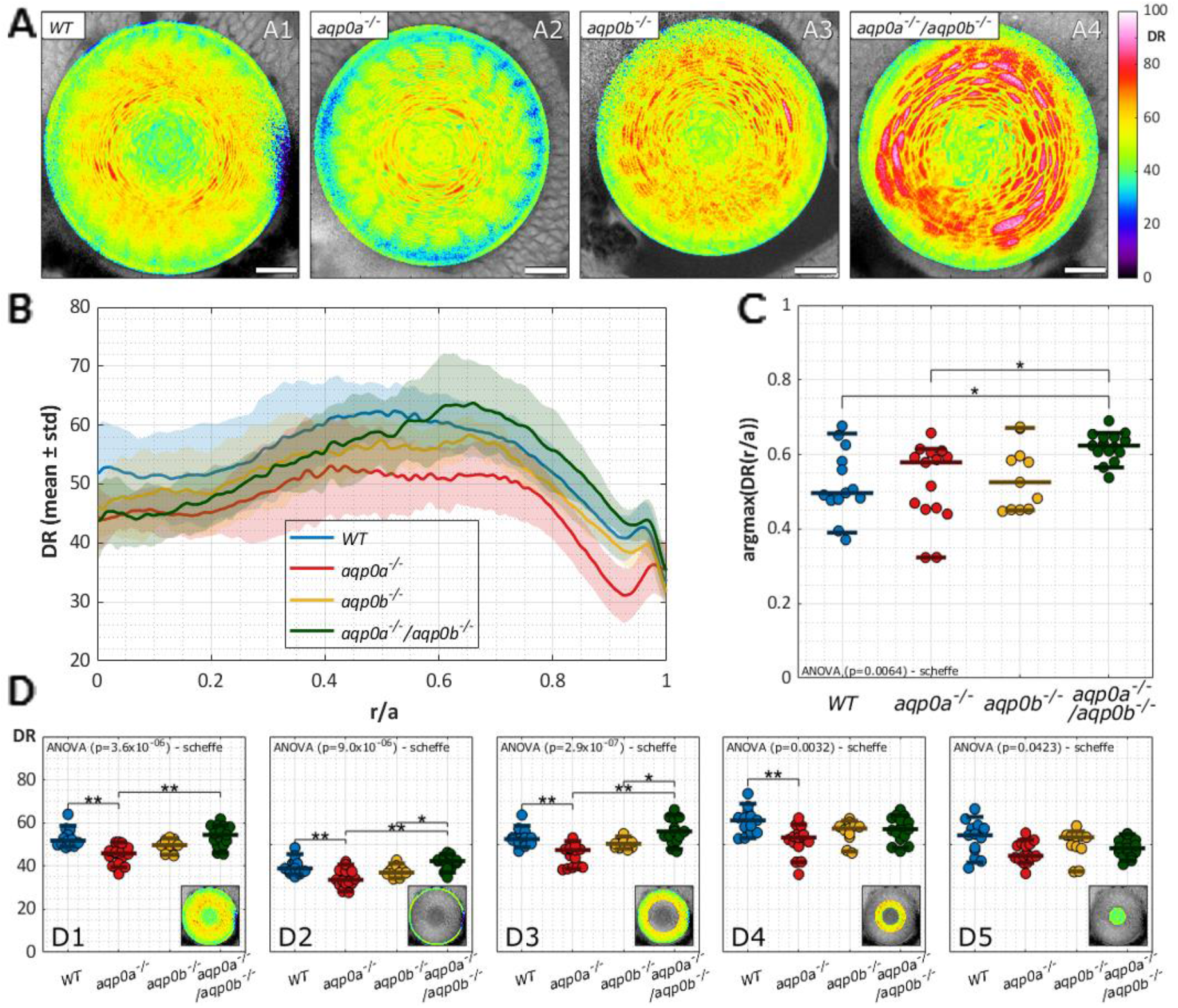
Disruption of dipolar relaxation distribution in Aqp0 mutant lenses. **(A)** Examples of dipolar relaxation (DR) images of lenses at 4 dpf of WT and mutants. Scale bars are 20 μm. **(B)** DR radial profiles (N=54). **(C)** Radial lens position of the maximal DR value in different mutants. **(D)** Mean DR of the whole equatorial lens plane, epithelium, outer cortex, inner cortex, and core of the lens as indicated by the insets. See Supplementary Table 3 for a summary of n numbers.

We next examined the lens poles (see Supplementary Table 1 for specific locations). At the anterior pole we have previously shown that *aqp0a*^*-/-*^ mutants show severe suture defects at older stages (21). ACDAN emission analysis in anterior lens planes revealed that *aqp0a*^*-/-*^ mutant lenses had lower DR (Figure 6A2) than other genotypes, which were all quite similar to one another (Figure 6A1, 3-4). This lower DR in *aqp0a*^*-/-*^ compared to the other genotypes was confirmed by comparison of the mean DR of the fiber cells (Figure 6B3), including the suture (Figure 6B4). In contrast, cells in the epithelium showed no difference (Figure 6B2, Supplementary Figure 11A). The maximum DR signal was closer to the center in *aqp0a*^*-/-*^*/aqp0b*^*-/-*^double mutant lenses than WT (Supplementary Figure 11B).

**Figure 6:**
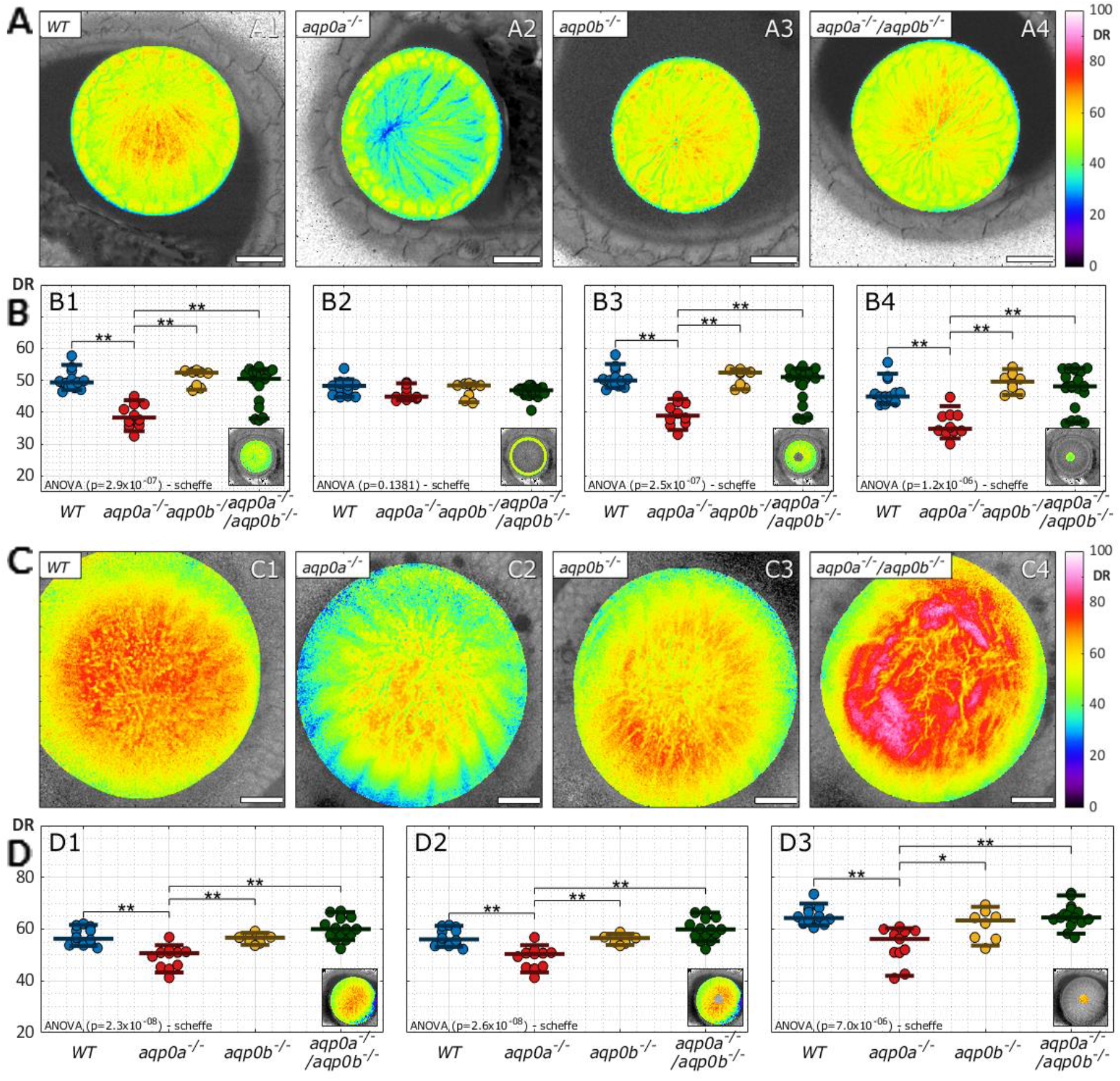
Dipolar relaxation in anterior and posterior planes of Aqp0 mutant lenses. Examples of dipolar relaxation (DR) images were taken at anterior planes (10 μm from the anterior pole) of lenses at 4 days post-fertilization. Scale bars are 20 μm. **(B)** Mean DR of the whole anterior lens plane **(B1)**, the epithelium **(B2)**, fiber cells (B3), and sutural regions **(B4)** as indicated by the insets (N=50). **(C)** Examples of DR images taken at posterior planes of lenses (75-85 μm from the anterior pole). Scale bars are 20μm. **(D)** Mean DR of the whole posterior pole **(D1)**, fiber cells **(D2)**, and sutural region **(D3)** as indicated by the insets (N=46). See Supplementary Table 3 for a summary of n numbers.

At the posterior pole, aqp0a-/-lenses also exhibited lower DR than other genotypes (Figure 6C2), while double *aqp0a*^*-/-*^*/aqp0b*^*-/-*^ mutant lenses had regions with higher DR that appeared as swollen cells (Figure 6C4). Cell morphology was severely disrupted, and these lenses lacked a clear convergence of a suture compared to the other genotypes (Figure 6C1-3). Statistically, *aqp0a*^*-/-*^ had a lower DR in the cortex and sutural regions of the posterior cortex compared to the other genotypes, while the double mutant was not statistically different (Figure 6D1-3), likely due to variability in the severity of the phenotype (See Supplementary Figure 10). The mean DR was relatively even around the sutures within ∼20 μm radius, with a dip at the center (Supplementary Figure 11C), which is likely why the maximum DR was very scattered around this value (Supplementary Figure 11D). Interestingly, the DR was overall lower in whole lenses, fiber cells, and sutures at the anterior pole than the posterior pole in all genotypes (Supplementary Figure 8).

Reconstructions of z-stacks in axial orientations confirmed the DR patterns and phenotypes observed in lenses imaged as single optical slices (Figure 7). Taken together, these results suggest that both zebrafish Aqp0s facilitate fluid efflux, disruption of which leads to lower DR and swollen fiber cells in the lens periphery, but that only Aqp0a facilitates influx, which is required to develop and maintain a higher DR throughout the lens cortex.

**Figure 7:**
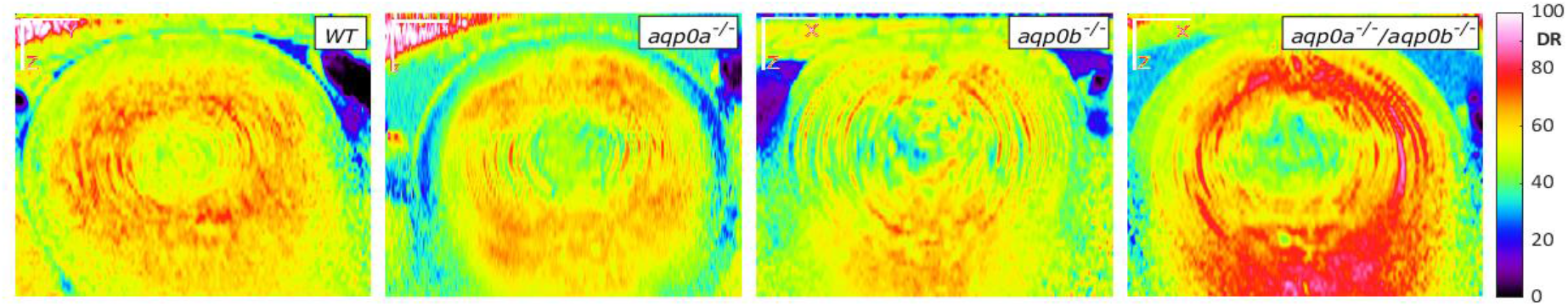
Axial orientation of DR signal in Aqp0 mutant lenses. Examples of dipolar relaxation (DR) axial slices through the center of the lens in *WT, aqp0a*^*-/-*^,*aqp0b*^*-/-*^ and *aqp0a*^*-/-*^*/aqp0b*^*-/-*^ double mutant lenses reconstructed from z-stacks. Anterior is oriented up. Scale bars are 20μm.

### Restoration of Aqp0 lacking water transport function fails to rescue macromolecular crowding defects in Aqp0-deficient lenses

To test whether it is the water transport property of the Aqp0s that is required for establishment and maintenance of lens water homeostasis, we employed WT and water-transport-dead aquaporin 0 DNA constructs and tested their ability to rescue *aqp0a*^*-/-*^ and *aqp0a*^*-/-*^*/aqp0b*^*-/-*^ mutant lens DR phenotypes. Aquaporin 0 from the killifish (*Fundulus heteroclitis*), MIPfun, was used to rescue the DR phenotype. Transiently expressed MIPfun has previously been able to rescue embryonic cataract due to knock-down of Aqp0a or Aqp0b, therefore, it is likely to encompasses functions of both zebrafish Aqp0s, while displaying similar, high water permeability of zebrafish Aqp0s compared to mammalian AQP0 (22, 36). Mosaics with strong expression of the transgenesis marker, mCherry, were selected for analysis (Figure 8B). The observed phenotypes varied due to variability in the mutant phenotypes’ penetrance, especially of the double *aqp0a*^*-/-*^*/aqp0b*^*-/-*^ mutant (see Supplementary Figure 10), and were further exacerbated by mosaicism of the rescue. Therefore, here we report the most consistent phenotypes with examples.

**Figure 8:**
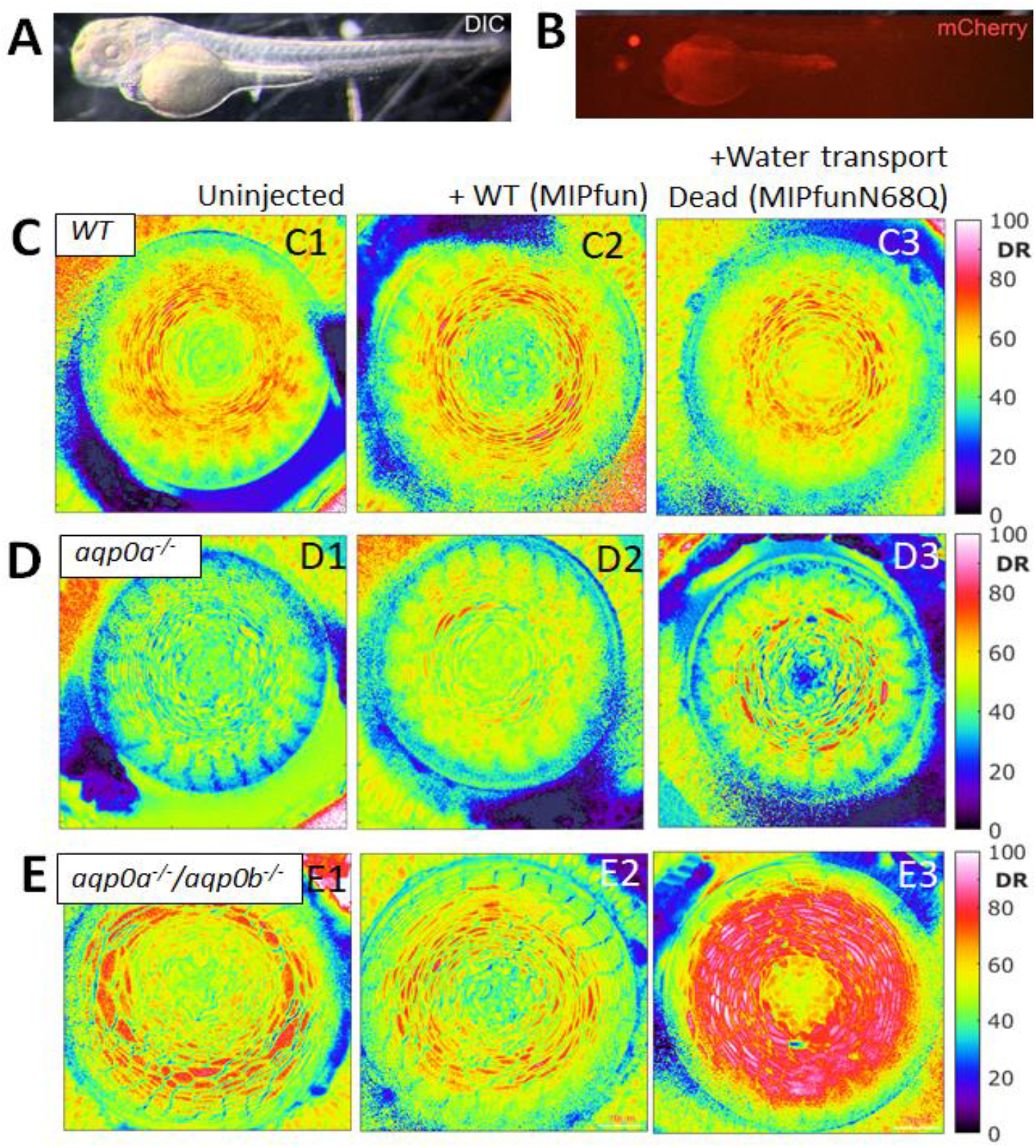
Water channel dead rescue did not restore water homeostasis in the lens. **(A)** Example of zebrafish injected with a rescue construct imaged under DIC illumination and strong expression of the transgenesis marker, mCherry, in the lens with some autofluorescence from the yolk. Examples of dipolar relaxation (DR) images of **(C)** WT, (D) *aqp0a*^*-/-*^, and (E) *aqp0a*^*-/-*^*/aqp0b*^*-/-*^double mutant 4 dpf lenses in equatorial orientation uninjected and injected with WT MIPfun (*Heteroclitisfundulus* aquaporin 0) rescue construct *Tg(HuβB1cry:MIPfun-IRES-mCherry)*, and water transport dead construct *and Tg(HuβB1cry:MIPfunN68Q-IRES-mCherry)*. Representative lenses of at least 4 experiments are shown.

Injection of either rescue construct into WT lenses did not affect the lens DR or morphology (Figure 8C2, 3). WT MIPfun rescued the low DR of *aqp0a*^*-/-*^ (Figure 7D2), and the high DR of swollen cells in double *aqp0a*^*-/-*^*/aqp0b*^*-/-*^ mutant was also less severe (Figure 7E2). Both of these transgenics appeared more like the uninjected WT lens (Figure 7C1) compared to uninjected mutant lenses (Figure 7D1, E1). The water-transport-dead mutant construct, MIPfunN68Q, failed to rescue the mutant phenotypes and made them more severe (Figure 7D3, E3). This fact confirms that the water transport function is essential for Aqp0a, and for Aqp0b when Aqp0a is also missing, in establishing and maintaining lens water homeostasis, and thus macromolecular crowding environment in the lens cortex.

## Discussion

To address macromolecular crowding in the zebrafish lens development and to test the requirements of Aqp0 on its homeostasis, we used hyperspectral imaging of the solvatochromic probe ACDAN as a nano-sensor able to measure water activity in living zebrafish lenses. Since water is the most abundant dipole active molecule within cells, interactions with macromolecules can be detected by changes in ACDAN fluorescence resulting in a continuum of water dipolar relaxation (DR) that ultimately gives a read-out of cellular water dynamics in the ACDAN nano-environment (29). Notice that ACDAN can relax only the few water molecules in close proximity (Angstrom range), and that water relaxation depends on water concentration (water/solute ratio) and the water activity. This last concept refers to the possibility of sensing water with different rotational times, which happens when water is interacting with molecules (confined water), and its relaxation is compromised when compared with bulk water (nanosecond vs. picosecond relaxation time, respectively)(32). Molecular crowding can modify both, the water/solute ratio and the water activity, and here we demonstrate that water DR, as measured with ACDAN fluorescence, provides an accurate measurement of crystallin protein crowding in solution (Figure 1).

The 2D and 3D imaging analysis tools developed in this work allowed for a consistent, partially-automated spatial and temporal study of water DR. This customized approach is crucial to quantify the regional lens water DR distribution enabling analysis and comparison of hundreds of *in vivo* optical lens slices. The intensity image of the ACDAN fluorescence (Supplementary Figure 6A, 9A) reveals no relevant information about the water DR distribution across the lens. However, transformation of the hyperspectral data into the phasor plot reveals a clear map of the water DR by using the reciprocity principle to generate the macromolecular crowding map at subcellular resolution in the lens (Supplementary Figure 6C, 9C). In the water DR image, our color scale represents the spectral shift identified at the phasor by the phase change (Figure 2B and C). Therefore, the water DR image (Figure 2C) highlights the power of this spectroscopic approach for *in vivo* lens imaging.

The first finding was that DR in the lens (regardless of the development stage, Figure 3) was lower than the most concentrated crystallin solution (Figure 1C). In absolute units, the lowest water DR obtained for the ACDAN in living lenses was ∼30 DR, which is much lower when compared to ∼95 DR found in concentrated crystallin solution (340 mg/ml). This result indicates that the water in the interior of the lens cells is more confined and is suggestive of a gel-like matrix compared with solutions (37, 38). This outcome is in line with a higher total concentration of different crystallins in the *in vivo* lens (up to 60% total mass) compared to what can be reached in solution due to crystallin aggregation (6). *In vivo* aggregation is largely prevented by chaperones in the lens (39). Furthermore, lens cell compaction, cytoskeletal and intermediate proteins, lipids, and many more macromolecular crowders required for normal lens refractive properties contribute to its unique macromolecular environment. Thus, within the lens nucleus water is strongly confined (low-activity), limiting the possibility of ACDAN to relax it, emphasizing the tight control of the macromolecular crowding for cellular and lens proper function.

The DR parameter from the spectral phasor of ACDAN hyperspectral imaging in the developing zebrafish lens reveals a progressively lower DR signal in the lens nucleus compared to the cortex, consistent with previous evidence of fiber cell compaction, concentration of crystallins (34), thereby increasing the protein to water ratio in the lens nucleus. Interestingly, ACDAN hyperspectral imaging reveals that water is handled differently during development at the anterior and posterior zebrafish lens sutures, likely the result of regional differences in water influx in early development. Notice that the variation in DR values found in the lens implies significant spatial and temporal changes in the macromolecular crowding compared with the water DR variation in response to the crystallin concentration in solution. Our analyses of *aqp0a-/-* and/or *aqp0b-/-* mutants unravels that while Aqp0a plays an essential and unique role in the regulation of water homeostasis, Aqp0b plays a secondary role that is revealed with the loss of both Aqp0s. This result is consistent with previous studies suggesting divergent functions for the two zebrafish Aqp0s, with Aqp0b more important in other functions in lens fiber cells, such as adhesion (21).

In addition to water, macromolecular crowding is affected by pH, ion concentration, and electrochemical gradients. In mature mammalian lenses pH decreases (40), intracellular Na^+^ (41) and Ca^2+^ (42) concentrations increase, hydrostatic pressure increases (43) and the plasma membrane depolarizes from ∼-70 to ∼-30 mV as a function of lens depth from the periphery to the center of the lens nucleus (Figure 9A)(8). Using ACDAN, we characterized the development of DR as lenses acquire these characteristics. Most striking is the decrease in DR in the lens nucleus with development, with the maximum DR shifting to the cortex (see Figure 3). This shift in the DR maximum is evident at 4 dpf (lens diameter ∼100-120 μm). Interestingly, at 3 dpf, the highest DR was measured in the center of the lens, similar to earlier stages, despite the lens nucleus having lost its organelles by 65 hours post-fertilization (35), and appearing tightly packed morphologically (24, 44). This high DR index at 3 dpf suggests that water has not been transported out of the nucleus and the macromolecular crowding of the lens nucleus is still ongoing and that the optics are immature.

**Figure 9:**
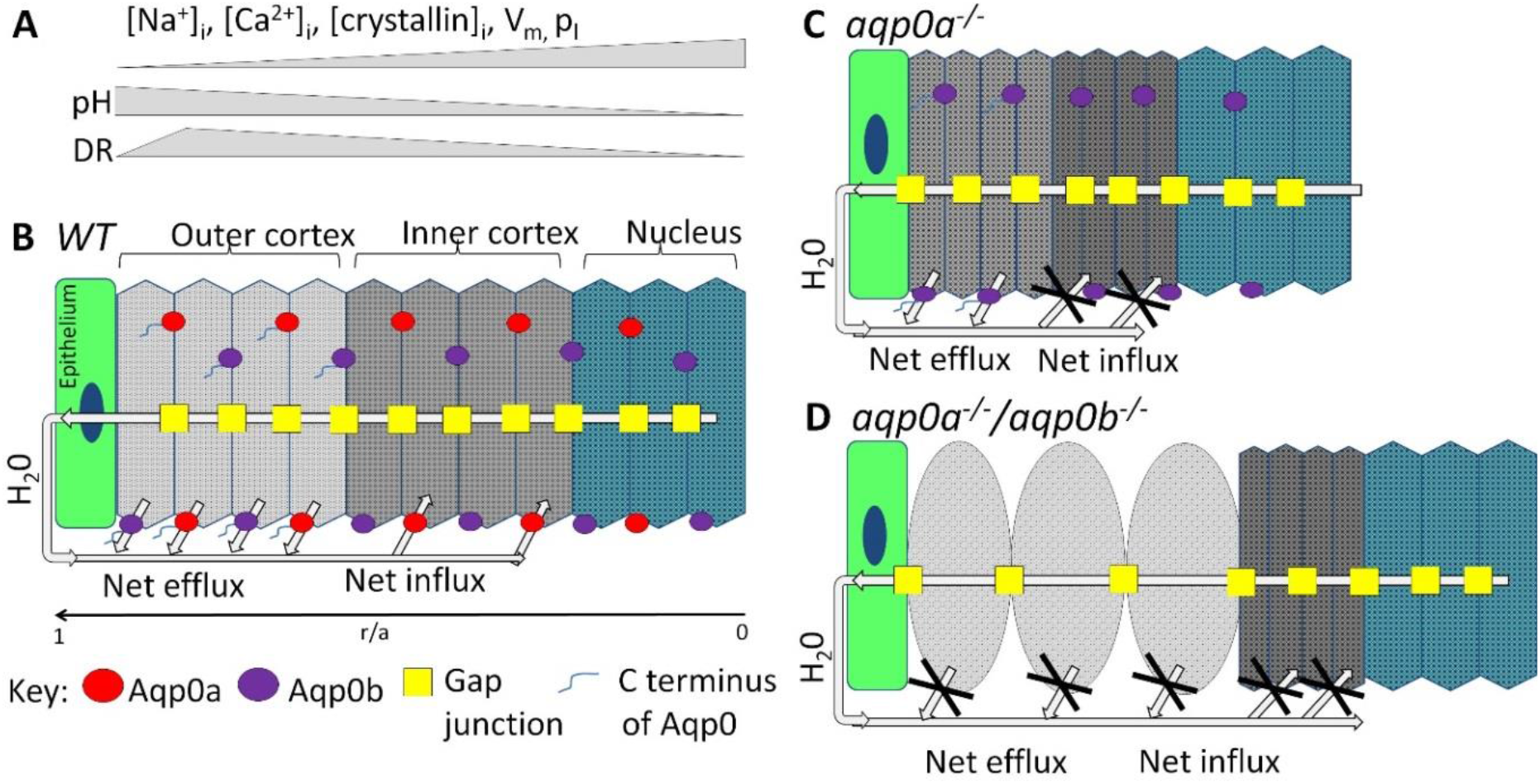
Model of the role of Aqp0a and Aqp0b in lens water transport *in vivo*. **(A)** Summary of physiological changes: intracellular Na^+^ ([Na^+^]_i_), Ca^2+^ ([Ca^2+^]_i_), crystallin concentrations ([crystallin]_i_), plasma membrane voltage potential (V_m_), lens hydrostatic pressure (p_i_) and intracellular pH are shown as relative changes by grey bars, that occur from the outer cortex to the nucleus in a mature lens correlating to lens regions in **(B)**. These changes determine its regional macromolecular crowding and thus dipolar relaxation (DR). DR peaks in the outer cortex in mature lenses, and decreases in the lens nucleus indicating lowest macromolecular crowding and highest water activity is in the outer lens cortex, and lowest water activity is in the highly crowded lens nucleus. **(B)** Diagram of a mature equatorial lens fiber cell stack from epithelium (r/a=1), to the center of the lens nucleus (r/a=0) with direction of H_2_0 flow shown based on previously published work. Aqp0a (red) and Aqp0b (purple) localize to both, broad and narrow fiber cells. Analysis of DR in *aqp0a*^*-/-*^**(C)** and *aqp0a*^*-/-*^*/aqp0b*^*-/-*^**(D)** lenses shows that both, Aqp0a and Aqp0b facilitate osmotic water efflux in the outer cortex, while only Aqp0a facilitates water influx into fiber cells in the inner cortex **(B). (C)** Loss of Aqp0a results in a net loss of water influx, but Aqp0b is still able to facilitate efflux, resulting in net loss of water leading to increased macromolecular crowding and lower DR in the outer and inner cortex. **(D)** In double *aqp0a*^*-/-*^*/aqp0b*^*-/-*^ mutants, both the H_2_0 influx and efflux pathways are disrupted. This water loss results in a more crowded environment in the inner cortex, and lower crowding in the swollen cells of the outer cortex.

It is interesting to discuss these results in the context of recent study on organelle degradation in the lens by PLAAT phospholipases (for phospholipase A/acyltransferase)(45). Morishita et al. demonstrated that Plaat 1 is crucial for organelles degradation; since without organelle degradation, transparency and required refractive properties of the lens cannot be achieved. Our decrease in water DR in the normal lens nucleus after 4 dpf correlates with the results of Morishita, where they found a dramatic spatial and temporal reorganization the lens macromolecular environment. Moreover, our *aqp0a*^*-/-*^ and double mutant *aqp0a*^*-/-*^*/aqp0b*^*-/-*^ results show that even with all correct PLAAT machinery resulting in specified organelle degradation, it is the macromolecular crowding and water homeostasis that ultimately dictate the optical properties of the lens. Future studies should focus on the change in lens DR transition from 3-4 dpf to investigate how it is affected by pH, ion concentration, or electrochemical gradients, and even changes in PLAAT machinery activity.

Hyperspectral imaging of ACDAN also revealed striking differences in the DR signal in the anterior and posterior regions of the lens. DR peaked in the posterior lens around 4-5 dpf in the cortex and suture, and then dropped at older stages (see Figure 4). The zebrafish lens germinal zone is further posterior at embryonic and larval stages (44) than in mammalian lenses. It is therefore likely that increased water influx is facilitated in this region to allow rapid growth and elongation of fiber cells, reflected by high DR. As lenses mature, the germinal zone shifts more anteriorly towards the center of the lens in the optical axis, correlated with a reduced DR at the posterior pole. Newly formed sutures tighten leading to low DR at both poles.

A net influx of ions and fluid in the inner cortex and net efflux in the outer cortex at the equator of the mammalian lens is crucial for the microcirculation system that delivers nutrients and removes waste from deeper lens tissue (reviewed by Donaldson et al)(10). Our DR analyses with ACDAN support this model and suggest that both zebrafish Aqp0s facilitate fluid efflux, but only Aqp0a facilitates influx (Figure 9). There is no evidence of extracellular space dilations, which would indicate that cell membranes have separated due to loss of adhesion. Therefore, it is unlikely that the loss of presumptive adhesive property of Aqp0b leads to the phenotype observed.

In this model derived from our work (Figure 9), loss of Aqp0a results in a reduction of water entering the lens, and thus the intracellular environment of the cortex is more crowded, resulting in lower DR values in all cortical regions. When both, Aqp0a and Aqp0b are missing, both fluid influx and efflux are disrupted, resulting in cell swelling in the periphery, particularly near the posterior pole, and shrinkage in the deeper cortex. This swelling is marked by high DR in the swollen cells, indicative of lower macromolecular crowding. Any role for Aqp0b in water influx appears to be dispensable and compensated by Aqp0a, as *aqp0b*^*-/-*^ lenses resemble WT. Anterior and posterior poles of *aqp0a*^*-/-*^ mutants have reduced DR compared to WT, indicating a reduced influx throughout the lens cortex (see Figure 6). Aqp0a water transport function near the poles correlates with sutures and lens nucleus’ centralization defects. This transport fails in *aqp0a*^*-/-*^ at older stages (21). In contrast to the cortex, our DR data suggest that Aqp0a and/or Aqp0b are not essential for the maintenance of water homeostasis in the zebrafish lens nucleus, at least at 4 dpf.

Alternatively, alteration in DR and cell swelling in double mutants could reflect a completely separate role for Aqp0b in another function in lens fiber cells, such as cell adhesion. However, since *aqp0b*^*-/-*^ mutants lenses look like WT, it is likely that the loss of its presumptive adhesive properties is compensated by other mechanisms, such as gap junctions. Our rescue experiments provide evidence to support the model that the water transport functions of Aqp0a and Aqp0b are required for proper macromolecular crowding, and by inference water homeostasis (see Figure 8). We show that WT MIPfun, which likely possesses functions of both zebrafish Aqp0s, reduces the severity of the DR reduction in aqp0a^-/-^ mutants, most likely by restoring the water influx. WT MIPfun also reduces the severity of cell swelling in double mutants, consistent with restoration of water influx and efflux. In contrast, mutant MIPfun constructs lacking water transport function (MIPfunN68Q) fail to rescue the DR phenotypes, and exacerbates its severity. The introduction of a non-functional form of Aqp0 could be more detrimental than a missing Aqp0, as AQP0 monomers have been shown to work cooperatively in a tetramer (46). Thus, the mutant MIPfunN68Q could disrupt the function of the native Aqp0s, exacerbating the phenotype. These data confirm the essential role of Aqp0a and Aqp0b as water channels for maintenance of fluid influx/efflux balance in the lens cortex, and thus overall lens homeostasis due to a tight macromolecular crowding tuning.

In conclusion, macromolecular crowding is essential in all living cells, and in this study we employed hyperspectral imaging of the nano-environment sensor, ACDAN, to describe the development of macromolecular crowding in the living zebrafish lens at a subcellular level. The combination of spectroscopy tools and imaging processing analysis enabled us to report the very high macromolecular crowding in the lens compared with a high concentration of crystalline in solution. Besides, we show that as lens optics develop, DR increases indicating increased macromolecular crowding in the lens nucleus from 2-4 dpf. We also show that *aqp0a*^*-/-*^ mutant lens cortex had reduced macromolecular crowding and cell swelling in double *aqp0a*^*-/-*^*/aqp0b*^*-/-*^ mutant lenses. These results indicate that both zebrafish Aqp0s facilitate fluid efflux in the lens cortex, but only Aqp0a facilitates influx in the living zebrafish lens. In the future, we will test the requirements of amino acids known to regulate Aqp0 water transport by external Ca^2+^ and pH on DR on water influx and efflux in the lens cortex. This study also provides tools and methods for studying water dynamics and macromolecular crowding mechanisms in other tissues in living organisms.

## Materials and Methods

### Zebrafish husbandry

The animal protocols used in this study adhered to the ARVO Statement for the Use of Animals in Ophthalmic and Vision Research and have been approved by the Institutional Animal Care and Use Committee of University of California, Irvine protocol #AUP-20-145. Zebrafish (AB strain) were raised and maintained under standard laboratory conditions (47), except methylene blue was excluded from the embryonic media (EM) as this yielded background fluorescence during hyperspectral imaging. The aqp0a-/- and/or aqp0b-/- mutants were generated as previously described (21). 0.003% 1-phenyl-2- thiourea (Sigma, St Louis, MO, P7629) was added to EM from 20-24 h postfertilization to prevent pigment formation. From 6 dpf, larvae were fed a diet of live rotifers (47).

### Rescue constructs

For rescue of mutant phenotypes, the Tol2 transposable element system (48) was used to stably integrate WT *Tg(HuβB1cry:MIPfun-IRES-mCherry)* or water-channel-dead *Tg(HuβB1cry:MIPfunN68Q-IRES-mCherry)* constructs of the *Heteroclitisfundulus*aquaproin 0 (MIPfun). A 200 bp region of the human βB1 crystallin promoter (49) was used to drive expression specifically in the lens, and only lenses strongly expressing the transgenesis marker (IRES-mCherry) were used to assess rescue of the phenotype. By using spectral phasors, we were able to extract ACDAN data without interference from mCherry (50), so it was selected as a transgenic marker. Previously, MIPfun had successfully rescued MO-knockdown-induced transient cataracts at 3 dpf of Aqp0a or Aqp0b, so it is thought to encompass properties of both zebrafish Aqp0s (23), MIPfunN68Q mutation results in an inactive water channel aquaporin (23), and so was used to test the requirement for water channel function to rescue DR phenotypes.

### Crystallin preparation

The Antarctic toothfish (Dissostichusmawsoni) γM8d crystallin (GenBank, DQ143983) sample was kindly provided by Dr. Jan Bierma from Dr. Rachel Martin’s lab. The recombinant proteins were grown, purified and stored as previously described (51). Dilution series from 0-340 mg/mL were made in buffer (10 mM phosphate pH 6.9, 50 mMNaCl, 0.05% NaN_3_). Concentration was measured by a Nanodrop at absorbance 280nm, and corrected by *ε*_*280*_=1.063 mL/mg at 1 nm.

### ACDAN staining

ACDAN (Toronto Research Chemicals, North York, ON-Canada, A168445) was dissolved in DMSO at 67 mM stock concentration, and added fresh to EM at a final concentration of 100 µM for overnight incubation of zebrafish prior to imaging. ACDAN was added at a final concentration of 5 µM 10 minutes prior to imaging of the crystallin preparation.

### Hyperspectral imaging

Embryos and larvae were anesthetized in EM with 0.0165% w/v tricaine (Sigma, St. Louis, MO, A5040) and mounted in 1% low melt agarose (Sigma, St Louis, MO, T9284) in 35 mm glass bottom microwell dishes (MatTek Corporation, Ashland, MA, P35G-1.5-14-C) with the eye against the coverslip, with the optical path perpendicular to the imaging plane (Supplementary Figure 1). Imaging planes were kept consistent between lenses of specific age as summarized (Supplementary Table 1).

Hyperspectral fluorescence images were acquired using a Zeiss LSM710 META microscope (Carl Zeiss, Jena GmbH) with a 40× water immersion objective 1.2 N.A. (Carl Zeiss, Jena GmbH). The microscope was coupled to a Ti:Sapphire laser (Spectra-Physics Mai Tai, Newport Beach, CA) which produces 80 femtosecond pulses with a repetition rate of 80 MHz. A two-photon wavelength of 780 nm was used for ACDAN excitation. The average laser power illuminating the sample was maintained at the mW level. Hyperspectral detection was performed with the Lambda Mode configuration of the Zeiss LSM710 META, which consists of a 32 channel GaAsP array photomultiplier tube. The hyperspectral range collected was from 416 to 728 nm; each of the 32 channels had a bandwidth of 9.7 nm. Image acquisition was performed with a frame size of 1024×1024 pixels, and a pixel size of 100 nm. Hyperspectral data was processed using a custom routine developed in MATLAB (The Mathworks, Inc., Boston, MA), described in the following sections.

### Spectral phasor of hyperspectral Image

The image processing pipeline we developed uniquely for analyzing spectral microscopy images of ACDAN emission in the zebrafish eye lenses is based on the spectral phasor transform. This integral transform obtains two quantities (named G and S) from the spectral intensity distribution at each pixel which are used to create the phasor plot of an image (52). The Cartesian coordinates (G,S) of the spectral phasor plot are defined by the following expressions:

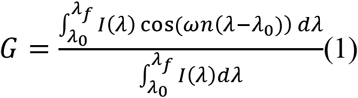

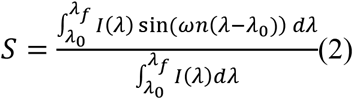

where*I(λ)* is the intensity as a function of wavelength at a particular pixel, measured in the interval (*λ*_*0*_*λ*_*f*_) that depends on the detector spectral range. The parameter n is the harmonic i.e. the number of cycles of the trigonometric function that are fit in the wavelength range by means of the angular frequency ω:

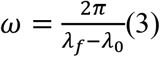

In practice one does not have a continuum of intensity values in the spectral direction, but rather a discrete number corresponding to the number of detectors that cover the spectral range. For computational purposes, the spectral phasor transform expressed as a discrete transform in terms of the spectral channel is (53):

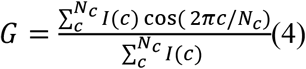

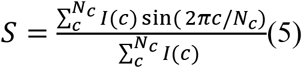

where now I(c) is the pixel intensity at channel c and N_C_ is the total number of channels. It is important that even if the number of spectral channels is small (in our case 32), the coordinates S and G are quasi continuous, due to the fact that the photon counts in each pixel and channel I(c) are high enough (∼102) to allow a wide range of values in the coordinates S and G.

The spectral phasor position of a particular pixel carries information about the spectral intensity profile of that pixel, allowing us to distinguish minute differences in the spectral emission. In polar coordinates, the angle carries the information regarding the spectral center of mass, and the radial direction carries information on the spectra broadness.

Most importantly though, spectral phasors follow rules of vector algebra, known as the linear combination of phasors (30, 54). This property refers to the additivity of components and allows the geometrical calculation of mixed pure environments. Pixels that contain a combination of two independent fluorescent species will appear on the phasor plot in a position that is a linear combination of the phasor positions of the two independent spectral species. The relative intensity fractions of the components determine the coefficients of the linear combination.

The other crucial property of the phasor plot is known as the reciprocity principle which refers to the fact that every point on the phasor plot corresponds to a pixel on the image and vice-versa, i.e. there is a bidirectional mapping between the image and the points. It is important to note that this operation is not a mathematical inversion; given the coordinates of a pixel in the phasor plot one cannot recover the photon spectral distribution of that pixel. This reciprocity maintained between the raw data and the phasor space representation allows us to select a region of interest in the phasor plot distribution and display the location of those pixels in the original image. An in-depth description of the properties of spectral phasor plots is given in references (32, 55).

The phasor transform applied to each pixel of an image produces a point in the phasor plot and all the pixels of an image together comprise the phasor plot. In the case of the particular range of wavelengths of our experiments and the range of our spectral detector array, this distribution was in a region between the first and second quadrant of the phasor plot. After plotting all the spectral images in the spectral phasor plot (a total of 420 images) we manually defined our region of interest in order to include all the points in the phasor plot in terms of a phase angle interval. This phase angle interval [*φ*_*0*_*φ*_*f*_] was chosen at [65°,115°] which approximately corresponds to the range [470 520]nm. This interval was then used to define our dipolar relaxation index as follows:

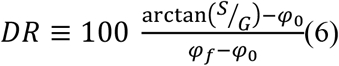

This quantity was then mapped to a particular lookup table (colormap) in order to color-code each pixel in the images according to the position of the pixel’s phasor transform in this interval (see Figure 2B-C). This color-coded image therefore now corresponds to a value in the interval [0,100], in turn corresponding to the aforementioned angular interval. It is this magnitude we refer to as the dipolar relaxation index (DR)(33). This DR definition does not require any prior knowledge of spectral characteristics of the sample.

A special consideration during the image processing steps regards saturated pixels in the images. This circumstance is uncommon since during the acquisition we ensured the use of a fraction of the dynamic range, but on rare occasions with a few outlier pixels this condition did not hold. Because in such pixels the spectral distribution is capped at some point, an error was introduced in the computation of the phasor transform. For this reason these pixels were marked before computing the phasor transform and the G and S phasor values for these pixels were interpolated a posteriori by averaging the neighboring pixels’ G and S values.

### Imaging processing routine for spatial/temporal study of the lens

In order to perform the spatial analysis, lenses were segmented from the background. Due to the fact that the lenses are not perfectly circular, and the challenge to align lenses perfectly to the imaging optical axis, and most importantly that the radial geometry of the cells conforming the lens does not in general match with the geometric center of the lens, the segmentation was performed in a semi-automatic way. For each image (420 total images), we manually marked six points around the edge of the lens, which were then used to automatically interpolate an arc joining them while forcing continuity in the arc and its derivative. The central point in the lens was used as the arc anchor, which had also been manually marked. This segmentation allowed tracing a total of 360 radii and obtaining the mean radial DR values in all directions. These curves were then interpolated to have equal numbers of points and were used to construct a Cartesian unfolding of the DR distribution of the lens (Supplementary Figure 2). From this Cartesian unfolded lens -one direction being the radius and the other the angle -the mean radial DR profile was obtained by projecting the mean DR in the angular direction (Figure 2D).

From the original circular geometry, a regional separation was applied to obtain the mean DR in each of the relevant regions of the lens. These regions were defined as annular bands taking into account the irregular geometry of the lens section, i.e. if the center of the lens is not the geometric center, in one direction the regional bands are tighter than in the other. For the equatorial plane a total of four regions were defined; epithelium (r/a<0.93), outer cortex (0.55<r/a<0.93), inner cortex (0.30<r/a<0.55) and core (r/a<0.30). For the anterior plane, the epithelium was manually segmented and both for the anterior and posterior planes, the suture was defined as the central circle of radius 10μm. The image processing experimental pipeline is represented in Supplementary Figure 3.

### Statistical analysis

When plotting results, we performed several fits to the data points. In such cases the coefficient of determination (R2; unity minus the sum of squared distances to the fit over the sum of squared distances to the mean) is provided to measure the goodness-of fit of the models used. Furthermore, a shaded area with a chosen confidence interval was also drawn in the background.

When comparing two independent distributions, the one-side Kolmogorov Smirnoff test for normality was performed in each of the two distributions. In the cases in which the test was passed, a Student t-test was used to statistically test if the data came from normal distributions with equal means. In the cases the normality test was not passed, the Wilkoxon rank sum test was used to statistically measure the chance that the two sets of points were drawn from distributions with equal medians.

When comparing more than two independent distributions, again the Kolmogorov Smirnoff test was used to test for normality of each distribution, and in the successful cases an ANOVA test was performed against the hypothesis that all groups are drawn from distribution with equal means. When the normality test failed the Kruskal Wallis test was used instead (56). In both cases, for further comparison of pairwise distributions for equal means, Scheffe’s procedure was chosen as it proved to be the most conservative.

## Supporting information

Supplementary Material: figure S1-11, Tables1-4, Supp Animations 1-6

## Acknowledgments

The authors are grateful to Prof. David Jameson, and Prof. Paul Donaldson for reading the manuscript and for their valuable suggestions. This work was supported in part by grants NIH P41-GM103540, NIH P50-GM076516, and NIH R01-EY05661. LM is supported by the Agencia Nacional de Investigación e Innovación (ANII) grant FCE_3_2018_1_149047, FOCEM -Fondo para la Convergencia Estructural del Mercosur (COF 03/11) and Chan Zuckerberg Initiative as Imaging Scientist. We thank Ines Gehring for zebrafish husbandry and assistance in generating and maintaining the Aqp0 mutant lines. We thank Drs. Jan Bierma and Rachel Martin for kindly providing the crystallin sample.

## Author contributions

I.V., A.V. L.M. designed experiments; I.V., A.V, B.T, L.M. carried out experiments; I.V. and A.V. analyzed the data; A.V. wrote software for data analysis, I.V., A.V., B.T. and L.M. wrote the manuscript, L.M., E.G. T.S and J.H. corrected the manuscript.

## Data and materials availability

Upon request, we will make the data available to other researchers. The spectral phasor analysis is part of a custom set of scripts in MATLAB. The full code is available upon request.

