## Supplementary Material: figure S1-11, Tables1-4, Supp Animations 1-6 for "*In vivo* macromolecular crowding is differentially modulated by Aquaporin 0 in zebrafish lens: insights from a nano-environment sensor and spectral imaging"

#### -Supplementary Materials-

**Irene Vorontsova[†,1,2,3], Alexander Vallmitjana[†,4], Belén Torrado[4], Thomas Schilling[2], James E. Hall[1], Enrico Gratton[4], Leonel S. Malacrida[¶,5,6]**

[1]Physiology and Biophysics, University of California, Irvine, Irvine, CA, USA.
[2]Developmental and Cell Biology, University of California, Irvine, Irvine, CA, USA.
[3]Neurobiology and Behavior, University of California, Irvine, Irvine, CA, USA.
[4]Biomedical Engineering, University of California, Irvine, Irvine, CA, USA.
[5]Departamento de Fisiopatologia, Hospital de Clínicas, Facultad de Medicina, Universidad de la República, Montevideo, Uruguay.
[6]Advanced Bioimaging Unit, Institut Pasteur of Montevideo and Universidad de la República, Montevideo, Uruguay.
[†]Equal contribution.

#### Supplementary Figures

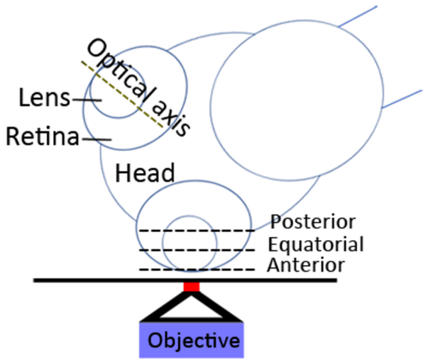


**Supplementary Figure 1**: **Zebrafish mounting.** Anesthetized embryos or larvae were mounted with the optical axis of the eye perpendicular to the imaging plane. The equatorial plane was selected to pass through the center of the lens nucleus, while anterior and posterior planes were obtained to capture the cross-section of the sutures at distances specified from the anterior pole in Supplementary Table 1.

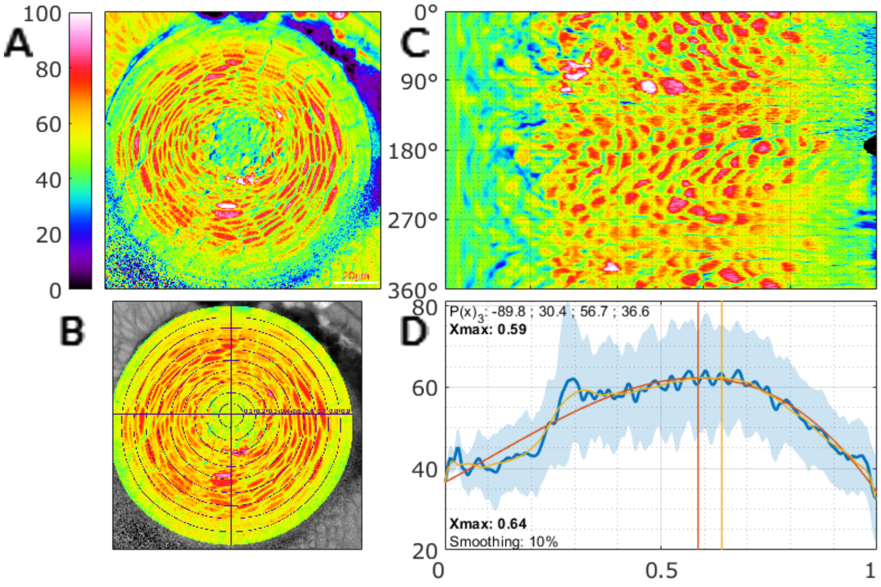


**Supplementary Figure 2**: **Obtaining radial dipolar relaxation profiles.** (A) Example of equatorial lens color-coded dipolar relaxation (DR) image. (B) Segmentation of lens edges and center, from which radial profiles are extracted. (C) Remapping of the radial profiles into an orthogonal space with angle in the vertical direction and radius in the horizontal. (D) Mean radial profile across all angles i.e. mean horizontal line in panel C, with the position of the maximum value marked (obtained in two ways; from a fitted 3rd degree polynomial and by smoothing).

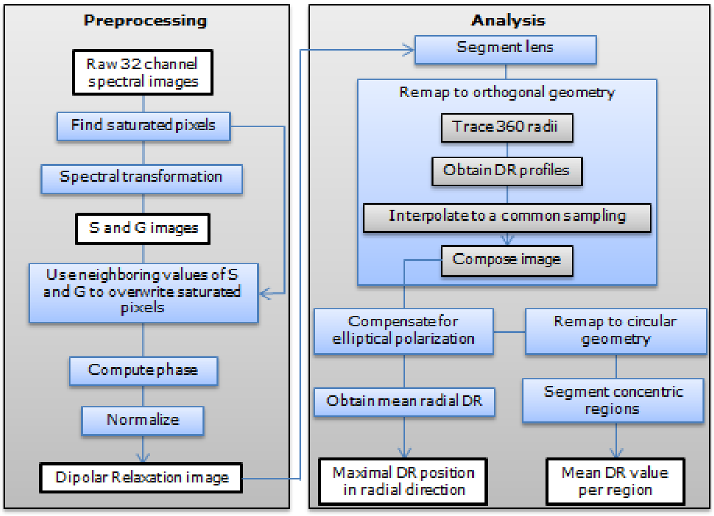


**Supplementary Figure 3**: **Image processing pipeline.**

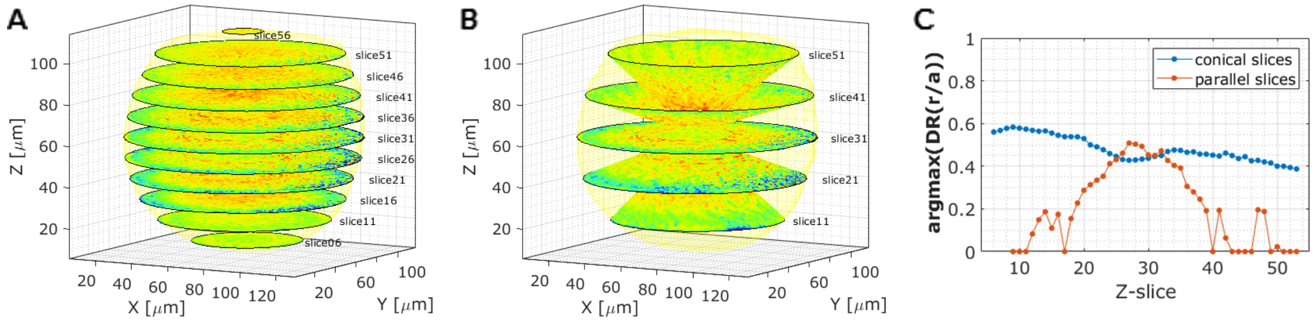


**Supplementary Figure 4**: **Dipolar relaxation has spherical symmetry opposed to cylindrical.** A z-stack of dipolar relaxation (DR) of a 4 dpf WT lens is used to segmented in 3D. (A) Selected slices are shown. (B) Selected conical sections are shown interpolated from the z-stacks. (C) The mean radial position of the maximum DR in each slice is plotted, both for the parallel and the conical slices.

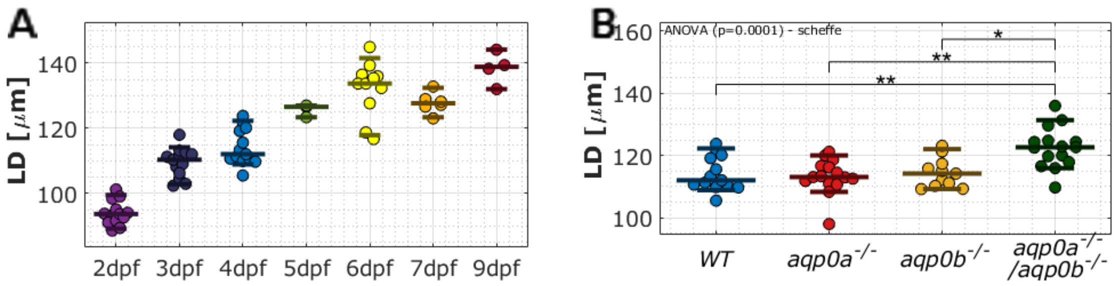


**Supplementary Figure 5**: **Lens size as a function of age and genotype.** (A) WT equatorial lens diameters (LD) increase as a function of their age in days post fertilization (dpf; N=63). However, variability of lens diameters within each group of dpf illustrates variability in fish growth, thus lens diameter is a better measure of developmental stage than age. (B) Equatorial lens diameter is wider in aqp0a$^{-/-}$/aqp0b$^{-/-}$ compared to other genotypes at 4 dpf (N=54).

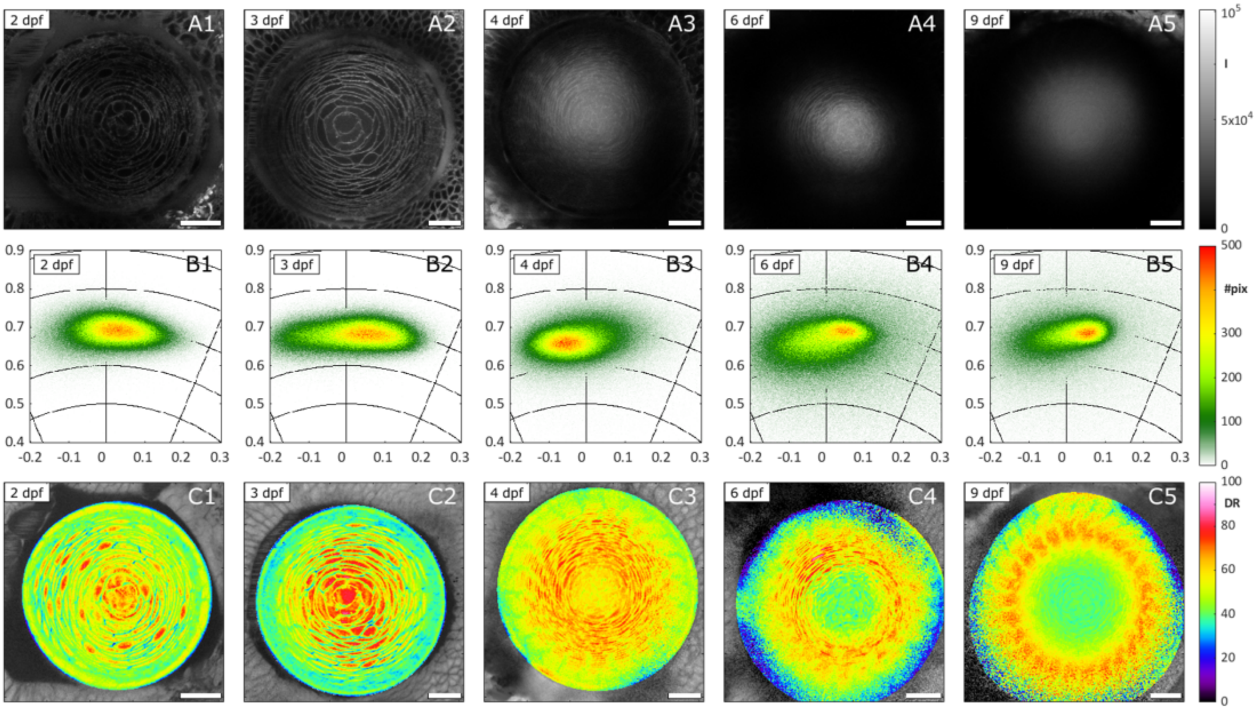


**Supplementary Figure 6**: **ACDAN intensity images to color-coded dipolar relaxation images with lens development.** (A) Intensity images of ACDAN in equatorial planes of WT lenses of specified days post fertilization (dpf), with their respective spectral phasor plots (B), and the segmented dipolar relaxation (DR) of the lens (C). These are the same lenses shown in figure 3 in the main body of the text.

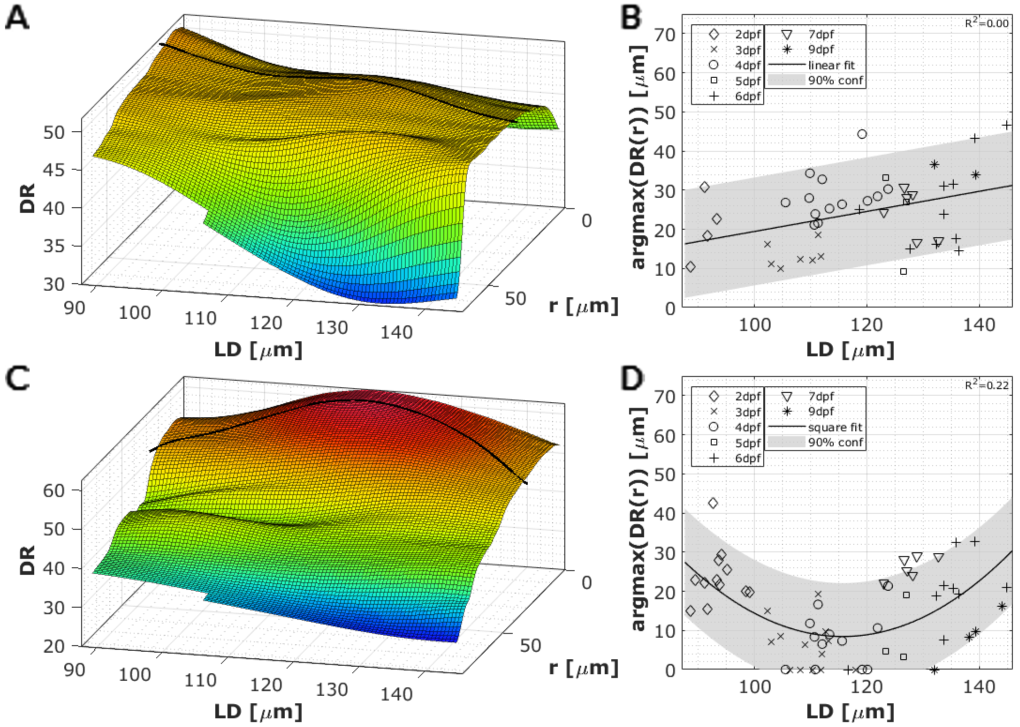


**Supplementary Figure 7**: **Mean and maximum dipolar relaxation distribution in anterior and posterior planes of zebrafish lenses during development.** A) Smoothed surface of the mean dipolar relaxation (DR) radial profile (radius not normalized, expressed in $\mu$m from lens center) as a function of lens diameter in anterior plane (N=45). (B) Radial lens position of the maximal DR value with development tends to slightly increase. A linear fit to the data and in turn represented on the surface in panel A. (C) Smoothed surface of the mean DR radial profile (radius not normalized, expressed in $\mu$m) as a function of lens diameter in posterior plane (N=59). (C) Radial lens position of the maximal DR value with development tends to slightly increase. A square polynomial is fit to the data and in turn represented on the surface in panel C.

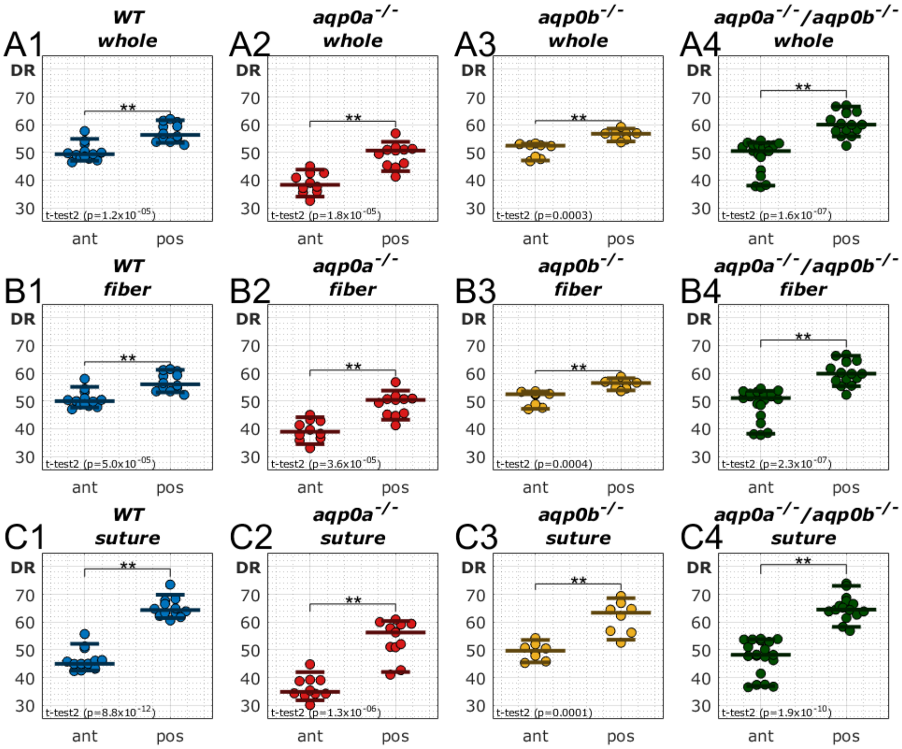


**Supplementary Figure 8**: **Mean dipolar relaxation is lower at the anterior compared to the posterior pole.** Mean dipolar relaxation (DR) from (A) whole anterior (ant) and posterior (pos) lens images, (B) fiber cells excluding the sutures and epithelium in case of anterior, and (C) sutural regions of the lenses were compared. Significance was denoted with ** at a significance level $p<0.005$. These data were extracted from Figure 5.

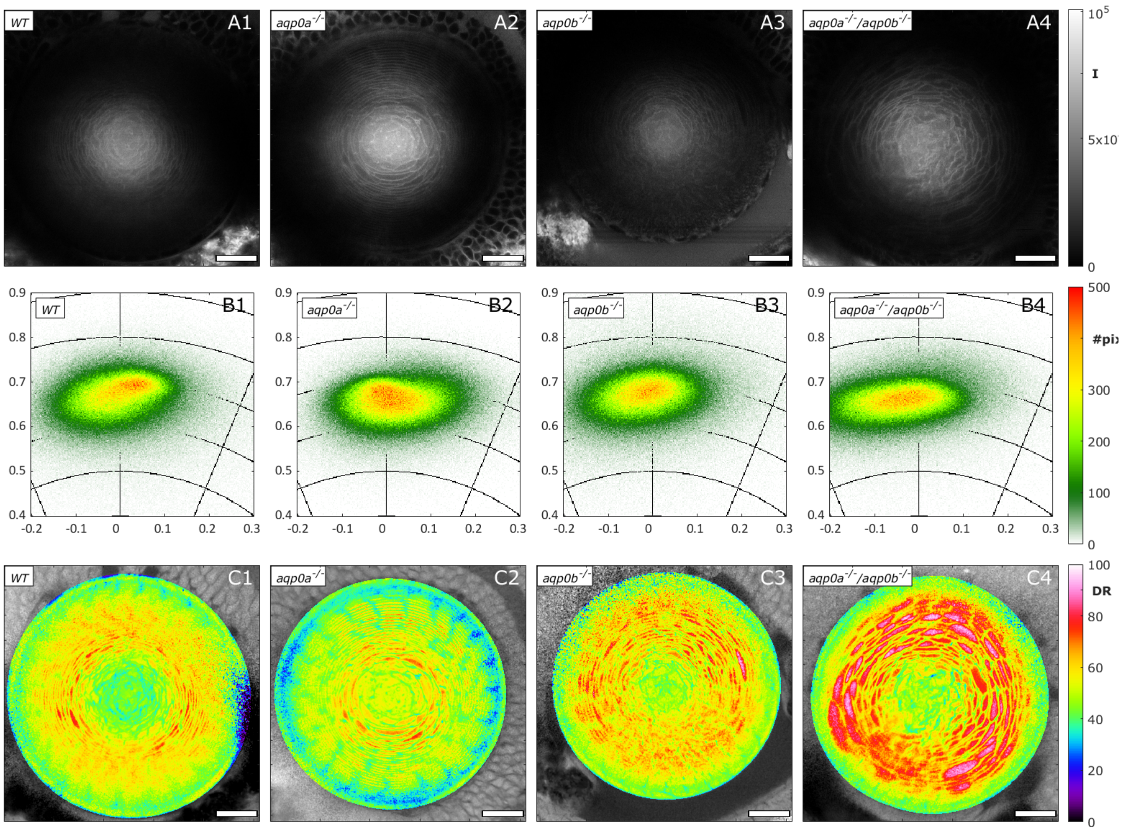


**Supplementary Figure 9**: **ACDAN intensity images to color-coded dipolar relaxation images in Aqp0 mutants.** (A) Intensity images of ACDAN in equatorial planes of lenses of specified genotype at 4 days post fertilization with their respective spectral phasor plots (B), and the segmented dipolar relaxation (DR) of the lens (C). These are the same lenses shown in figure 5 in the main body of the text.

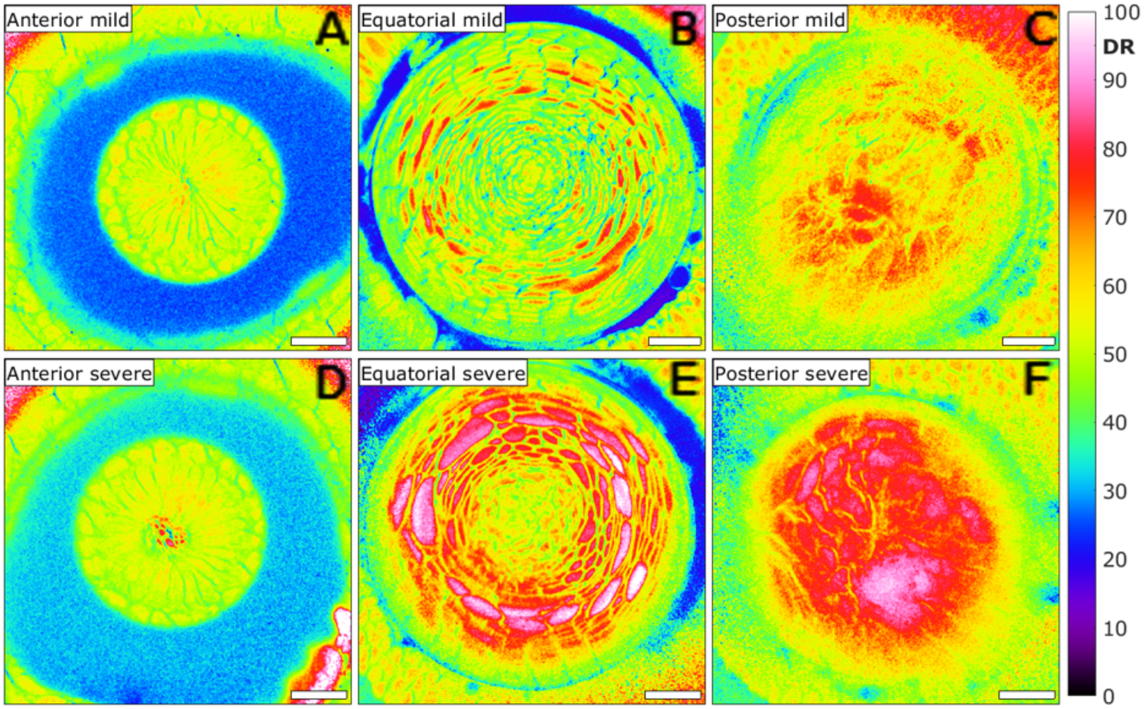


**Supplementary Figure 10**: **Dipolar relaxation phenotype variability in double aqp0a$^{-/-}$/aqp0b$^{-/-}$ mutants.** Dipolar relaxation (DR) images of double aqp0a$^{-/-}$/aqp0b$^{-/-}$ mutants revealed 13/19 lenses had WT-like anterior sutures (anterior mild, A), compared to 6/19 lenses with high DR anterior sutural regions (anterior severe, D). In the equatorial and posterior planes 8/15 had milder phenotypes (B, C), and 7/15 had severe phenotypes (E, F). Scale bars are 20$\mu$m.

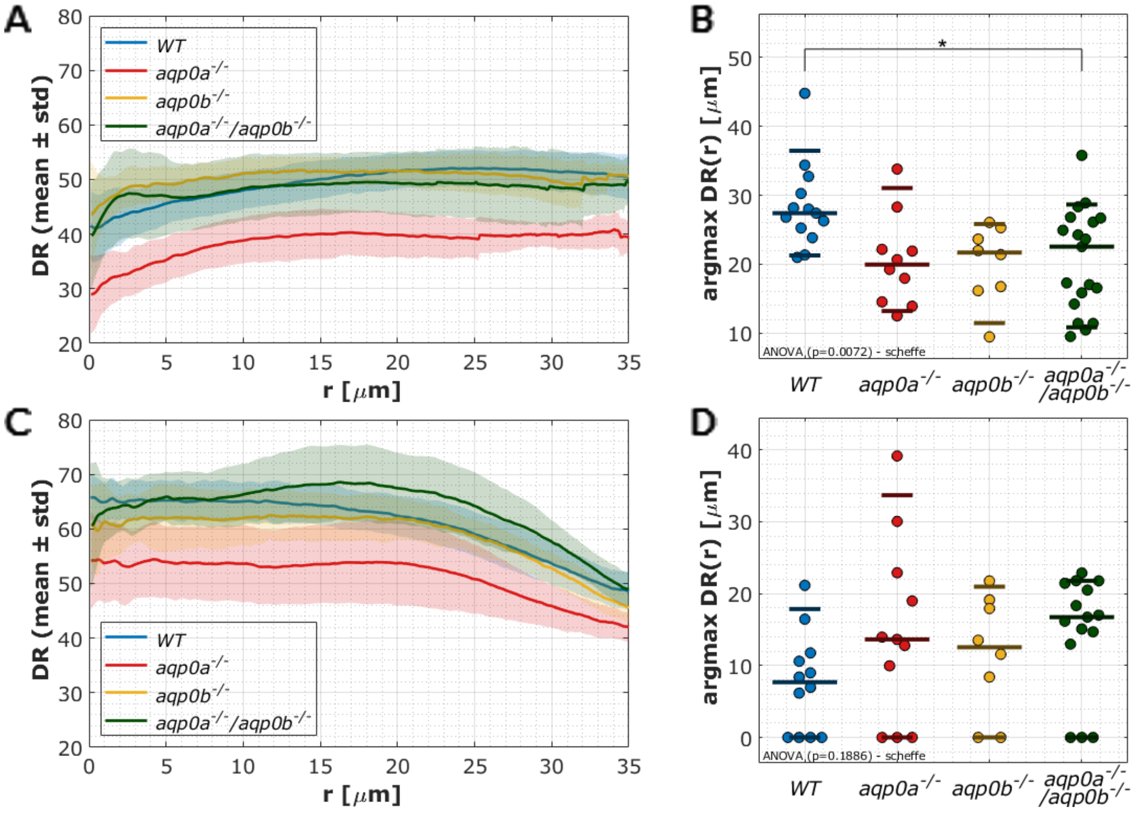


**Supplementary Figure 11**: **Mean and maximum dipolar relaxation distribution in anterior and posterior planes of Aqp0 mutant lenses at 4 dpf.** (A) Dipolar relaxation (DR) radial profiles as a function of distance from the suture in anterior planes of Aqp0 mutant and WT lenses (N=50). (B) Radial lens position of the maximal DR value in the anterior plane. (C) DR radial profiles as a function of distance from the suture in posterior planes (N=46). (D) Radial lens position of the maximal DR value in the posterior plane.

### Supplementary Tables

| Age | Distance imaged from the anterior pole ($\mu$m) | | |
|---|---|---|---|
| | Anterior plane | Equatorial plane | Posterior plane |
| 2 dpf | 5 | 45 | 85 |
| 3 dpf | 10 | 45-55 | 85 |
| 4 dpf | 10 | 45-55 | 85 |
| 5 dpf | 10 | 55 | 100-110 |
| 6 dpf | 10 | 55 | 140 |
| 7 dpf | 10 | 65 | 140 |
| 9 dpf | 10 | 65 | 140 |

**Supplementary Table 1**: Summary of distances from the anterior pole of acquisition of anterior, equatorial and posterior planes of lenses of different ages in days post fertilization (dpf).

| Plane | 2 dpf | 3 dpf | 4 dpf | 5 dpf | 6 dpf | 7 dpf | 9 dpf | Total |
|---|---|---|---|---|---|---|---|---|
| Equatorial | 13 | 13 | 13 | 3 | 11 | 6 | 4 | 63 |
| Anterior | 4 | 7 | 13 | 3 | 10 | 6 | 2 | 45 |
| Posterior | 12 | 13 | 12 | 3 | 9 | 6 | 4 | 59 |

**Supplementary Table 2**: Number of spectral lenses/images analyzed for each age in days post fertilization (dpf) and plane.

| Plane | WT | aqp0a$^{-/-}$ | aqp0b$^{-/-}$ | aqp0a$^{-/-}$ /aqp0b$^{-/-}$ | Total |
|---|---|---|---|---|---|
| Equatorial | 13 | 15 | 11 | 15 | 54 |
| Anterior | 13 | 10 | 8 | 19 | 50 |
| Posterior | 12 | 11 | 8 | 15 | 46 |

**Supplementary Table 3**: Number of lenses/spectral images analyzed in each plane.

| Genotype | Equatorial | Anterior | Posterior |
|---|---|---|---|
| WT | 13 | 13 | 12 |
| WT+ MIPfun | 7 | 7 | 7 |
| WT + MIPfunN68Q | 11 | 10 | 11 |
| aqp0a$^{-/-}$ | 15 | 10 | 11 |
| aqp0a$^{-/-}$ + MIPfun | 3 | 2 | 2 |
| aqp0a$^{-/-}$ +MIPfunN68Q | 6 | 6 | 6 |
| aqp0a$^{-/-}$/aqp0b$^{-/-}$ | 15 | 19 | 15 |
| aqp0a$^{-/-}$/aqp0b$^{-/-}$ + MIPfun | 5 | 5 | 5 |
| aqp0a$^{-/-}$/aqp0b$^{-/-}$ + MIPfunN68Q | 4 | 4 | 4 |
| Total | 85 | 82 | 79 |

**Supplementary Table 4**: Number of lenses/spectral images analyzed for rescue experiments.

### Supplementary Animations

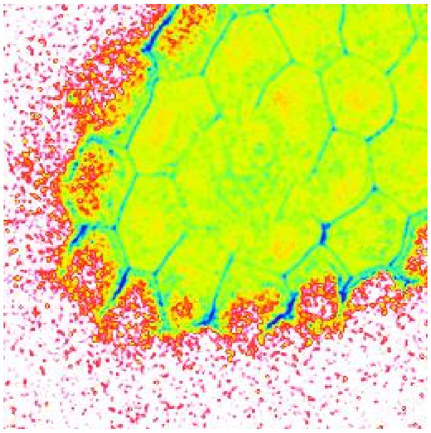

**Supplementary Animation 1**: **Cross-sections of the dipolar relaxation for a 2 dpf WT lens.** Individual frames of a z-stack animated through the optical axis (slices parallel to equatorial plane). Scale bars are 20$\mu$m.

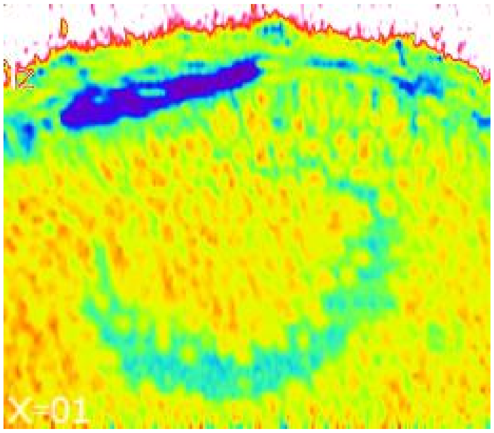


**Supplementary Animation 2**: **Cross-sections of the dipolar relaxation for a 2 dpf WT lens.** Interpolated frames from of a z-stack animated perpendicular to the optical axis (slices are parallel to the axial orientation of the lens). Scale bars are 20$\mu$m.

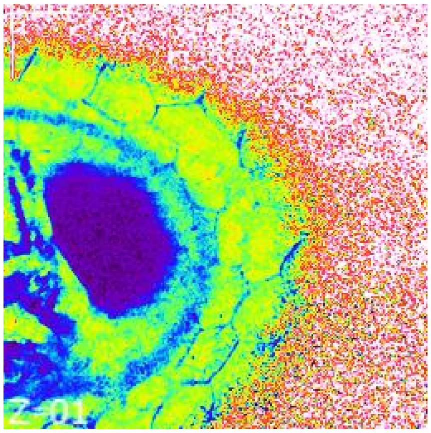


**Supplementary Animation 3**: **Cross-sections of the dipolar relaxation for a 3 dpf WT lens.** Individual frames of a z-stack animated through the optical axis (slices parallel to equatorial plane). Scale bars are 20$\mu$m.

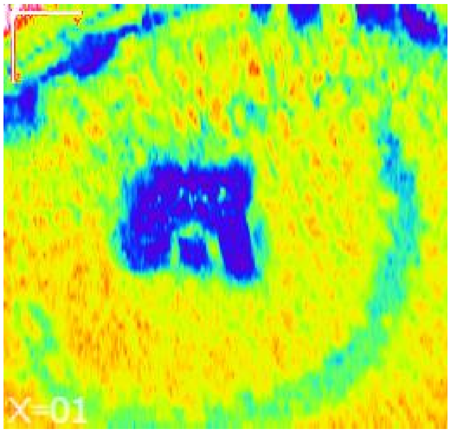


**Supplementary Animation 4**: **Cross-sections of the dipolar relaxation for a 3 dpf WT lens.** Interpolated frames from of a z-stack animated perpendicular to the optical axis (slices are parallel to the axial orientation of the lens). Scale bars are 20$\mu$m.

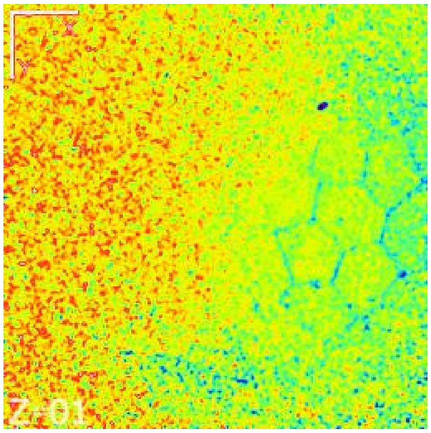


**Supplementary Animation 5**: **Cross-sections of the dipolar relaxation for a 4 dpf WT lens.** Individual frames of a z-stack animated through the optical axis (slices parallel to equatorial plane). Scale bars are 20$\mu$m.

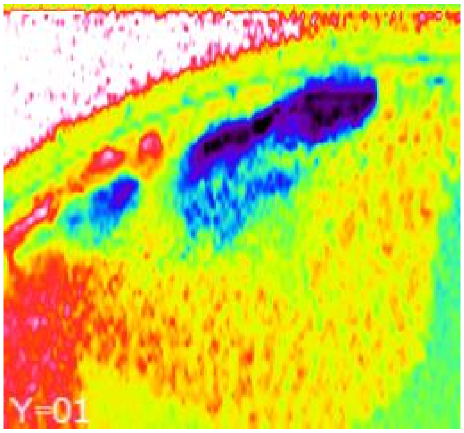


**Supplementary Animation 6**: **Cross-sections of the dipolar relaxation for a 4 dpf WT lens.** Interpolated frames from of a z-stack animated perpendicular to the optical axis (slices are parallel to the axial orientation of the lens). Scale bars are 20$\mu$m.
